# SRSF3 is a key regulator of epicardial formation

**DOI:** 10.1101/2021.11.25.470003

**Authors:** Irina-Elena Lupu, Susann Bruche, Anob M. Chakrabarti, Ian R. McCracken, Tamara Carsana, Andia N. Redpath, Nicola Smart

## Abstract

The epicardium is a fundamental regulator of cardiac development and regeneration, functioning to secrete essential growth factors and to produce epicardium-derived cells (EPDCs) that contribute most coronary mural cells and cardiac fibroblasts. The molecular mechanisms controlling epicardial formation have not been fully elucidated. In this study, we found that the RNA-binding protein SRSF3 is highly expressed in the embryonic proepicardium and epicardial layer. Deletion of *Srsf3* from the murine proepicardium led to proliferative arrest, which prevented proper epicardial formation. Induction of *Srsf3* deletion after the proepicardial stage resulted in impaired epicardial proliferation and EPDC formation by E13.5. Single-cell RNA-sequencing showed SRSF3-depleted epicardial cells were eliminated, however, the surviving non-recombined cells became hyperproliferative and, remarkably, compensated for the early deficit, via a mechanism that involved *Srsf3* up-regulation This unexpected finding attests the importance of SRSF3 in controlling epicardial proliferation, and highlights the significant confounding effect of mosaic recombination on embryonic phenotyping. Mapping the SRSF3–RNA interaction network by endogenous irCLIP identified binding to major cell cycle regulators, such as *Ccnd1* and *Map4k4*, with both splicing and non-splicing roles. This research defines SRSF3 as a key regulator of epicardial cell proliferation.

## Introduction

Understanding cardiac morphogenesis is essential for designing improved therapies to treat heart disease. The outer layer of the heart, the epicardium, plays a major role in cardiac formation and repair, shown to promote cardiomyocyte proliferation and vessel growth in developing embryos and in species that regenerate their hearts, through provision of growth factors and perivascular support^1–8^. For non-regenerative species, maximising the therapeutic potential of the epicardium depends upon augmenting the restricted reactivation and proliferation that occur endogenously^8^. The epicardial layer originates from the proepicardial organ (PEO), a transient embryonic structure located at the venous pole of the developing heart^9^. After epicardial formation is complete, around embryonic day (E) 11.5 in mouse, some epicardial cells undergo epithelial-to-mesenchymal transition (EMT), leading to the formation of epicardium-derived cells (EPDCs) that contribute most of the vascular smooth muscle cells and cardiac fibroblasts for the developing heart^10–14^.

A key process in epicardial formation and function is cellular proliferation. PEO cluster emergence in zebrafish was shown to be dependent on tension generated through regionalised proliferation of mesodermal progenitors^15^. Epicardial EMT also appears to require cell division, with orientation of the mitotic spindle dictating whether cells remain on the surface or invade the myocardium^16^. Studies in which epicardial proliferation rate was increased by Cyclin D1 overexpression^16^ or deletion of neurofibromin1 (NF1), a negative regulator of Ras^17^, resulted in increased formation of EPDCs.

The RNA-binding protein Serine/arginine-rich splicing factor 3 (SRSF3) is the smallest member of the serine/arginine rich (SR) family of proteins, master regulators of RNA metabolism in the cell, involved in multiple aspects of RNA processing, such as mRNA transcription, splicing, export, stability and translation^18–26^. SRSF3 is characterised as an oncogene due to its role in promoting cellular proliferation and migration in various cancers^22, 27–33^. Recently, SRSF3 was shown to be required for cardiomyocyte proliferation during development, with conditional deletion of SRSF3 from this lineage resulting in embryonic lethality mid-gestation^34^. SRSF3 has been implicated in multiple developmental transitions^35–38^, but the role of SRSF3 in the epicardium has not been previously investigated.

Here, we show that SRSF3 is expressed ubiquitously in the heart during embryonic development, with the highest expression levels in the PEO and in the epicardium until E12.5. To address the role of SRSF3 in the epicardium, we generated two SRSF3 conditional knockout mice. Deletion of SRSF3 using the Tg(Gata5-Cre) line resulted in impaired epicardial layer formation, due to decreased proliferation of founder cells in the PEO, and embryonic lethality at E12.5. Epicardial-specific deletion later in development, using the inducible Wt1^CreERT2^ line, resulted in a less severe phenotype characterised by impaired coronary vasculature formation, reduced cardiac compaction and myocardial hypoxia, due to defective epicardial proliferation and EMT. However, mosaic recombination allowed non-targeted epicardial cells to hyperproliferate, paradoxically up-regulating *Srsf3*, to drive a robust compensatory mechanism and virtually rescue the phenotype by late embryonic stages. To delineate the molecular mechanisms through which SRSF3 controls epicardial proliferation, we used single-cell RNA sequencing (scRNA-seq), to define the transcriptional changes that result from epicardial *Srsf3* depletion, and individual nucleotide resolution infrared cross-linking and immunoprecipitation (irCLIP), to map SRSF3-RNA binding sites across the epicardial transcriptome. These analyses confirmed control of mitotic cell cycle as a primary function of SRSF3 in the epicardium and demonstrated direct binding to transcripts encoding key regulators of proliferation, such as Cyclin D1, and senescence, including MAP4K4. These data reveal an essential role for SRSF3 in the proliferation of (pro)epicardial cells, and in the epicardial processes that underpin cardiac morphogenesis.

## Results

### Characterisation of SRSF3 expression during development

The expression pattern of SRSF3 at key stages of epicardial activity was investigated by immunohistochemistry (IHC) of mouse embryo and heart cryosections: including at E9.5, when epicardial progenitors first arise within the PEO; at E11.5, when the epicardium is most proliferative and has formed a layer^16, 39^; and at E15.5 when the epicardium downregulates key marker genes^39^ and progresses towards quiescence^10, 16^. The highest levels of SRSF3 were found in the PEO and in the early epicardium (E11.5), after which SRSF3 expression started to decline (Fig.1A, B, Supplementary Fig. 1A) until postnatal stages, when SRSF3 levels remained low (Supplementary Fig. 1B). Notably, reduction in SRSF3 expression levels coincided with the downregulation of Wilms’ Tumor 1 (WT1) in the epicardium (Fig. 1A), a key transcriptional regulator that marks the active epicardial state^8^. Strong SRSF3 expression was observed in E11.5 epicardial explant cultures (Fig. 1C), where outgrowth is governed by epicardial cell proliferation.

**Figure 1.**
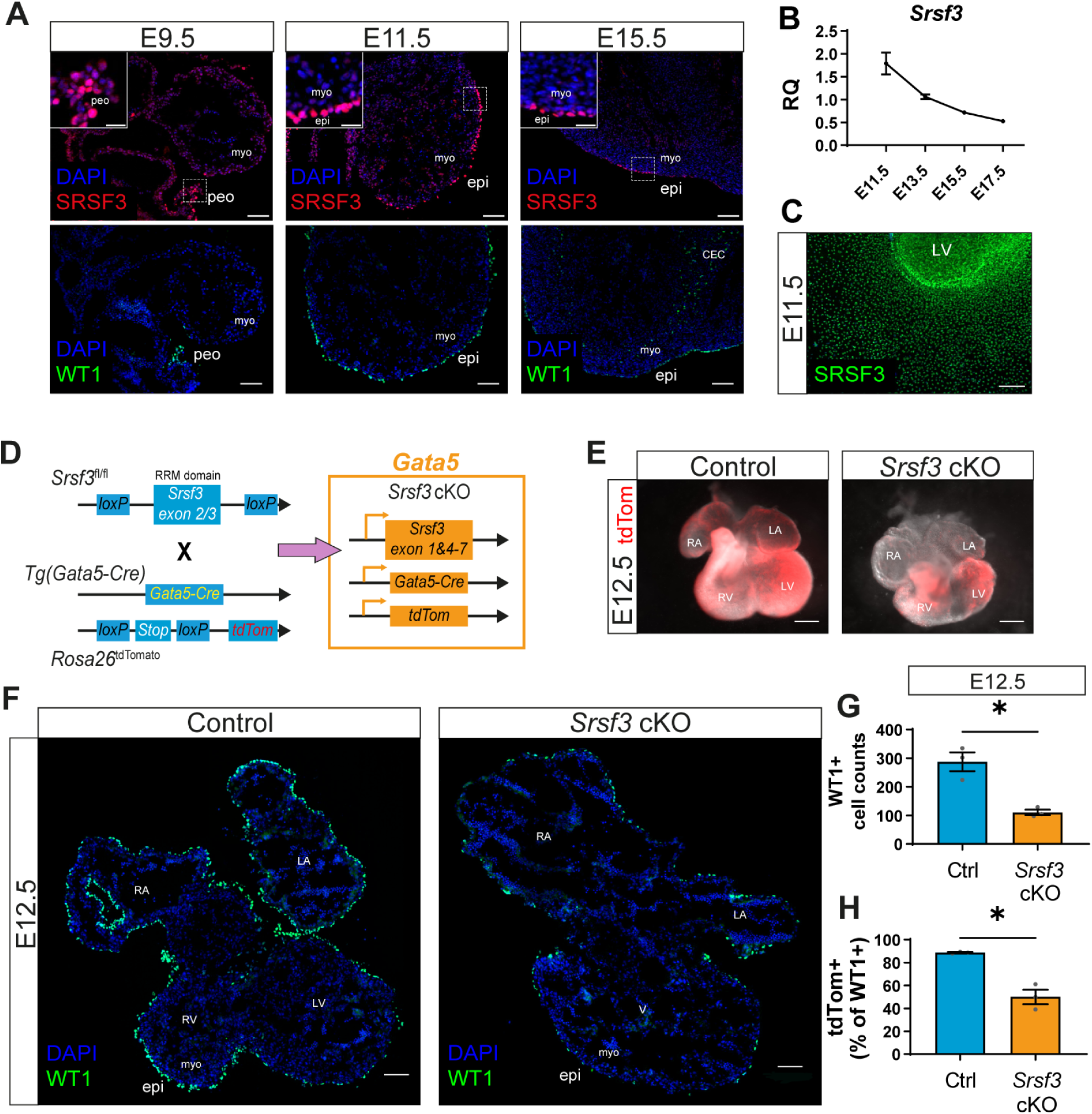
SRSF3 is expressed in the developing heart and is required for epicardium formation. (A) Immunofluorescence showing expression of SRSF3 and WT1 in embryonic mouse hearts at E9.5, E11.5 and E15.5. Scale bar, 100μm. Images representative of n > 4 embryos. (B) qRT-PCR analysis of *Srsf3* transcript expression in whole heart lysates at E11.5, E13.5, E15.5 and E17.5; RQ, relative quantification. Values normalized to *Actb*. Error bars indicate mean ± SEM (n = 5). (C) Immunofluorescence showing expression of SRSF3 in an E11.5 epicardial explant. Scale bar, 200μm. Image representative of n = 8 embryos. (D) Generation of a conditional SRSF3 knock-out mouse (*Srsf3* cKO) with lineage tracing capacity; targeting the epicardial lineage (Tg(Gata5-Cre)) and reported by tdTomato (Rosa26^tdTomato^). (E) Fluorescence stereo microscope images of control and *Srsf3* cKO embryonic hearts at E12.5. Epicardial lineage reported by tdTomato fluorescence. Scale bar, 500μm. Images representative of n = 3 embryos. (F) Immunofluorescence showing expression of WT1 in control and *Srsf3* cKO embryonic hearts at E12.5. Scale bar, 100μm. Images representative of n = 4 embryos. Quantification of (G) WT1+ cells and (H) percentage tdTomato+ epicardial lineage cells as a proportion of total WT1+ cells. Error bars indicate mean ± SEM (n = 3). Unpaired t-test with Welch’s correction (*) *P* <0.05. *myo; myocardium, peo; proepicardium, epi; epicardium, CEC; coronary endothelial cell, LV; left ventricle. RV; right ventricle, V; ventricles, RA; right atrium, LA; left atrium*.

### SRSF3 is required for epicardial formation and cardiomyocyte survival

To investigate the role of SRSF3 in the PEO, Tg(Gata5-Cre);Rosa26^TdTomato^ mice^40, 41^ were crossed with Srsf3^fl/fl^ mice^34^ in order to deplete SRSF3 in epicardial progenitor cells, and concomitantly label them to track their migration and fate (defined as *Srsf3* cKO; Fig.1D). Tg(Gata5-Cre) uses a chick enhancer of *Gata5* that becomes active from E9.25 and drives Cre recombinase expression in the PEO^40^, septum transversum and a subset of cardiomyocytes^42^. *Srsf3* cKO displayed embryonic lethality from E12.5, with no live embryos recovered beyond this stage (Supplementary Table 1). E12.5 *Srsf3* cKO embryos presented gross morphological abnormalities of the heart, including hypoplastic ventricles and dilated atria (Fig. 1E). Since the ventricles were smaller, cell death and proliferation were investigated by IHC. There was increased cell death and decreased proliferation in *Srsf3* cKO hearts, as indicated by cleaved-caspase 3 (CC3) and phospho-histone H3 (PHH3), respectively (Supplementary Fig. 1C-F). All the cells expressing CC3 were positive for sarcomeric-α-actinin (s-α-actinin), suggesting that SRSF3 is only required for cardiomyocyte survival (Supplementary Fig. 1C). There were fewer PHH3/tdTomato positive cells, both cardiomyocytes and epicardial cells, in *Srsf3* cKO hearts, compared with control hearts (Supplementary Fig. 1E, F), suggesting that SRSF3 is required for the proliferation of both cell types. The requirement for SRSF3 to enable cardiomyocyte proliferation in the developing embryo has been reported^34^, however, the involvement of SRSF3 in epicardial proliferation has not been addressed.

To investigate the impact of SRSF3 depletion on epicardial formation, immunostaining for WT1 was performed on E12.5 heart sections (Fig. 1F, Supplementary Fig. 1G). Fewer WT1 positive cells were detected on the surface of *Srsf3* cKO hearts compared to controls (Fig. 1F, G). It is important to note that ∼90% of epicardial cells were tdTomato labelled in E12.5 control hearts, as Tg(Gata5-Cre) was not active in all epicardial progenitor cells. The proportion of epicardial cells positive for tdTomato was significantly decreased in *Srsf3* cKO compared to controls (Fig. 1H, Supplementary Fig. 1G), suggesting that non-targeted cells are disproportionately either more proliferative or more likely to migrate onto the heart, compared with those in which *Srsf3* was successfully depleted.

### SRSF3 depletion in the PEO results in impaired proliferation and migration of epicardial progenitor cells

The impaired epicardial formation phenotype was more closely investigated in ventricular epicardial explants from E11.5 *Srsf3* cKO embryos, which demonstrated reduced outgrowth (Fig. 2A, B) and a lower proportion of tdTomato positive cells, compared with control (Fig. 2C). To assess if epicardial formation is disrupted from the outset, we checked for the presence of epicardial cells on the surface of the heart at E10.5, the stage when proepicardial progenitor cells complete their migration. Few WT1 positive epicardial cells were detected on E10.5 *Srsf3* cKO hearts, in contrast to the extensive coverage attained in controls (Fig. 2D, Supplementary Fig. 2A). This prompted us to determine whether the formation of the proepicardium itself was impaired by loss of *Srsf3*. Indeed, we observed fewer epicardial progenitor cells in the PEO of *Srsf3* cKO embryos at E9.5 compared to controls, with mostly non-targeted, tdTomato negative cells present (white arrows; Fig. 2E). This was likely a consequence of impaired proliferation, as demonstrated by diminished Ki67 expression (Supplementary Fig. 2B). To confirm the proliferative defect, and more accurately quantify progenitor expansion, PEO explants were cultured from control and *Srsf3* cKO embryos (Fig. 2F - H, Supplementary Fig. 2C). As expected, *Srsf3*-depleted PEO explants were significantly smaller than controls (mean 1166 versus 3654 cells per explant; Fig. 2F, Supplementary Fig. 2C) and consisted of 56% fewer Ki67 positive cells (Fig. 2G, H), further supporting the notion that SRSF3 is required for proliferation of epicardial precursor cells.

**Figure 2.**
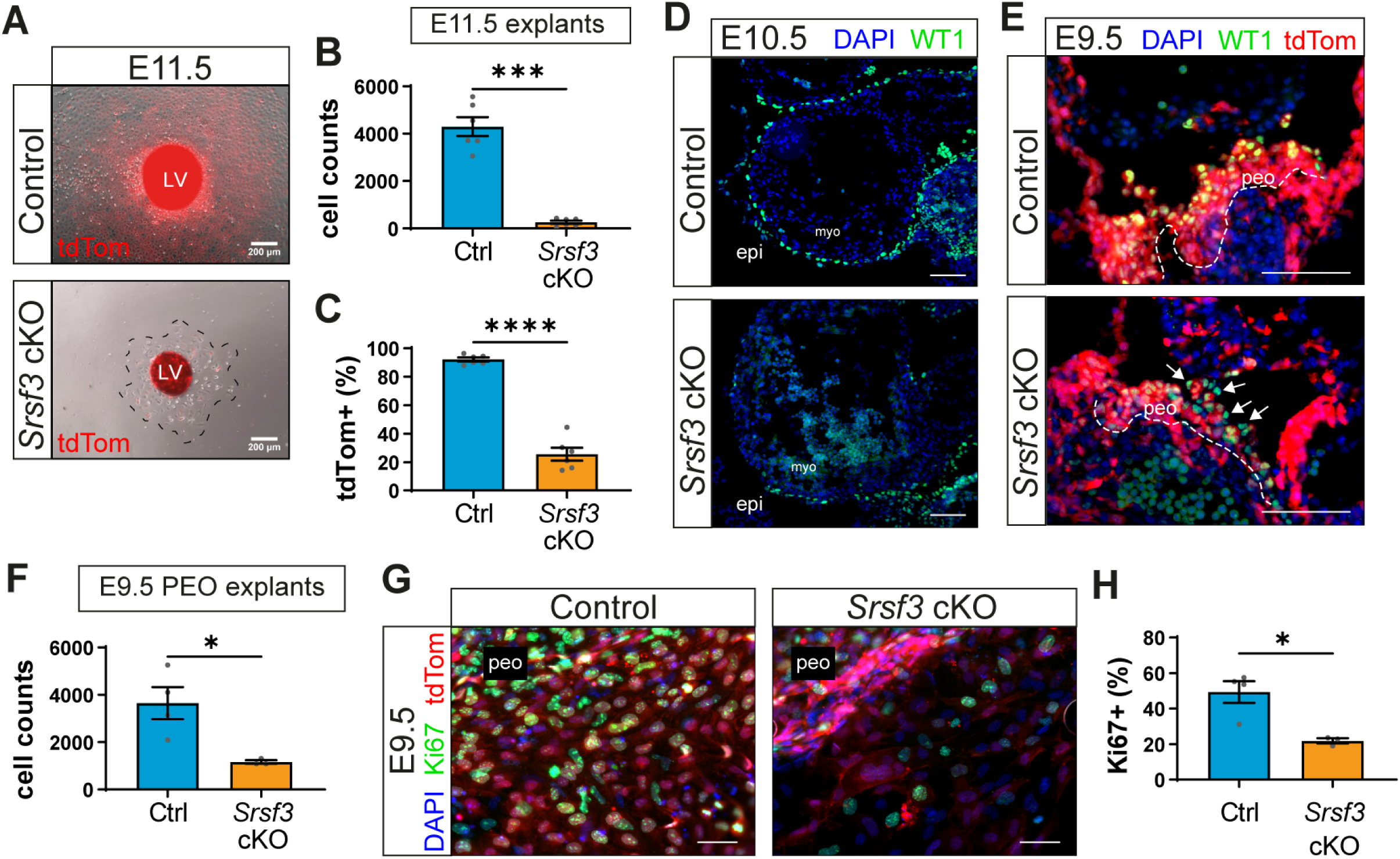
Impaired proliferation of epicardial progenitor cells in *Srsf3* cKO hearts. (A) Brightfield and tdTomato fluorescence images of control and *Srsf3* cKO E11.5 ventricular explants (3 days’ culture). Scale bar, 200μm. Quantification of (B) epicardial cells and (C) percentage of tdTomato+ cells. Error bars indicate mean ± SEM (n = 6). Unpaired t-test with Welch’s correction (***) *P* <0.001; (****) *P* <0.0001. (D) Immunofluorescence images showing WT1+ cells on the surface of control and *Srsf3* cKO embryonic mouse hearts at E10.5. Scale bar, 100μm. Images representative of n = 3 embryos. (E) Immunofluorescence showing WT1+ epicardial progenitor cells situated in the PEO (outlined by dashed line) of control and *Srsf3* cKO embryos at E9.5. White arrows highlight non-targeted cells. Scale bar, 100μm. Images representative of n = 5 embryos. (F) Quantification of (pro)epicardial cells from control and *Srsf3* cKO E9.5 PEO explants (day 3 culture). Error bars indicate mean ± SEM (n = 3 - 4). Unpaired t-test with Welch’s correction (*) *P* <0.05. (G) Immunofluorescence images and corresponding quantification of (H) percentage Ki67+ (pro)epicardial cells in control and *Srsf3* cKO E9.5 PEO explants (day 3 culture). Scale bar, 50μm. Error bars indicate mean ± SEM (n = 4 control, 3 *Srsf3* cKO embryos). Unpaired t-test with Welch’s correction (*) *P* <0.05. *myo; myocardium, peo; proepicardium, epi; epicardium, LV; left ventricle*.

### SRSF3 depletion in the epicardium leads to impaired proliferation and enhanced cell death in epicardial cells

To bypass the proepicardial defects and investigate SRSF3 function exclusively in the epicardium, we used the inducible Wt1^CreERT2^ line^43, 44^, with induction limited to E9.5 - E11.5 to avoid significant targeting of coronary endothelial cells^39^. This strategy was adopted to generate an inducible *Srsf3* KO (*Srsf3* iKO), with capacity to trace and temporally target the epicardial lineage (Fig. 3A). Embryonic hearts were initially evaluated at E13.5 when the epicardium is fully formed and EMT is underway. Despite having a largely intact epicardium, *Srsf3* iKO hearts exhibited a disrupted morphology and a marked reduction in the number of lineage-traced EPDCs that had invaded the myocardium (Fig. 3B). Moreover, embryos presented a severe non-compaction phenotype at this stage (Fig. 3C). To determine whether the EMT and compaction defects may be underpinned by impaired proliferation, given the *Srsf3* cKO phenotype, we assessed epicardial cell proliferation. Epicardial explants cultured from E13.5 *Srsf3* iKO hearts revealed a mosaic loss of SRSF3, yet, as expected, they contained fewer Ki67 positive cells (Fig. 3D, E), due to loss of proliferation where *Srsf3* was successfully deleted (Fig 3F, G; mean 37.2% versus 1.3%, in SRSF3+ and Srsf3-depleted cells, respectively). Impaired EMT and diminished emergence of EPDCs at E13.5 is consistent with the requirement for cell division to drive this process^16^, further supporting the role of SRSF3 in epicardial proliferation. Moreover, defective EMT and compaction coincided with impaired sprouting of vessels from the sinus venosus in *Srsf3* iKO hearts (Fig. 3H, I), which is also in keeping with the role of the epicardium in promoting coronary vessel growth^45^.

**Figure 3.**
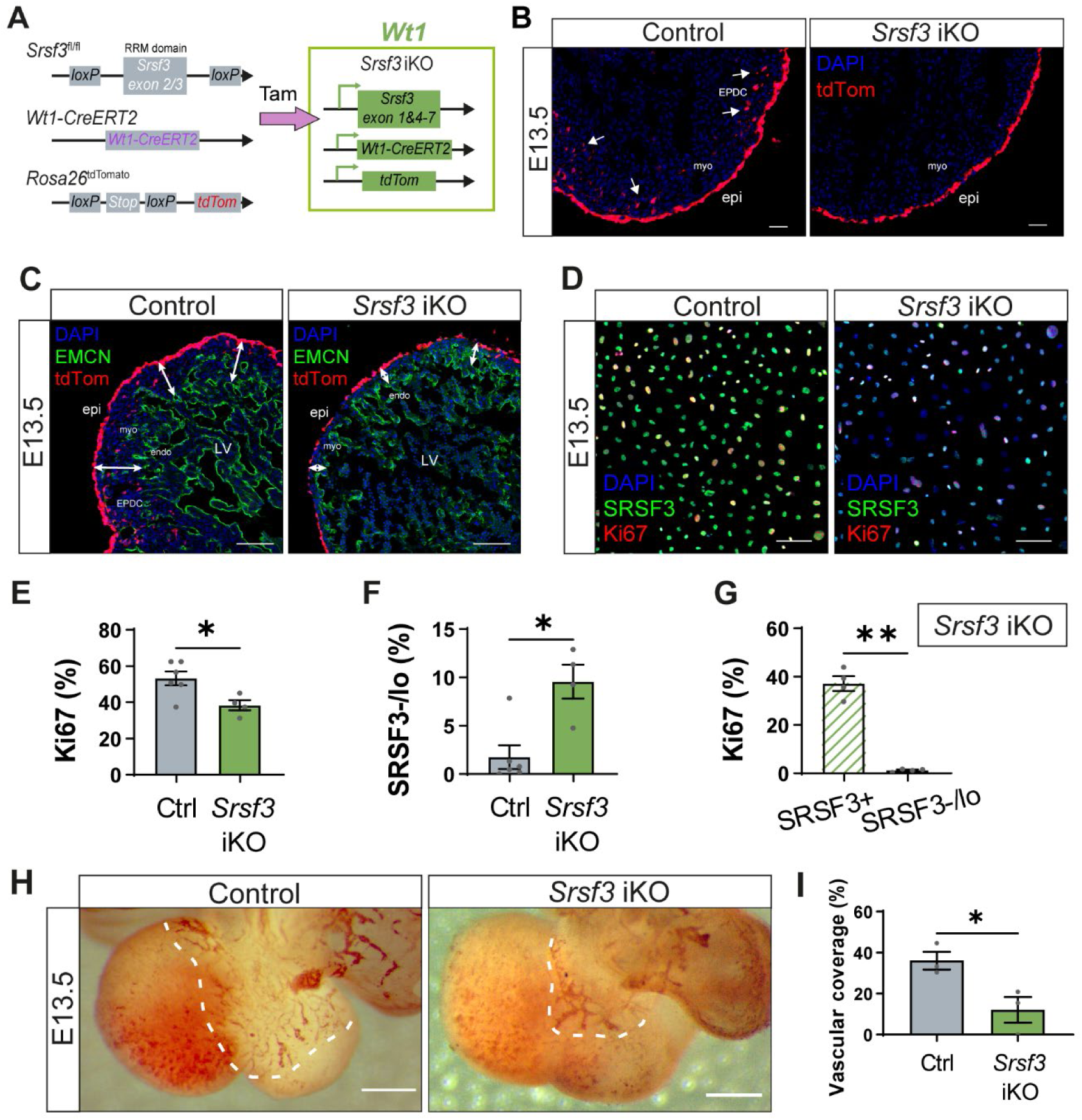
Impaired proliferation and function of epicardial cells in *Srsf3* iKO hearts. (A) Generation of an inducible SRSF3 knock-out mouse (*Srsf3* iKO) with lineage tracing capacity; restricted to the epicardial lineage (*Wt1*-CreERT2) and reported by tdTomato (Rosa26^tdTomato^). Tamoxifen was administered at E9.5/E11.5 (late-stage induction) (B) Fluorescence images showing epicardial lineage cells (tdTomato+) in control and *Srsf3* iKO embryonic mouse hearts at E13.5. White arrows highlight invasion of epicardium-derived cells (EPDC) into the myocardium. Scale bar, 50μm. Images representative of n = 6 embryos. (C) Immunofluorescence showing the endocardium and vascular network (EMCN+) in control and Srsf3 iKO embryonic mouse hearts at E13.5. tdTomato expression demarcates the epicardium lineage. White arrows highlight myocardial wall thickness. Scale bar, 200μm. Images representative of n = 4 embryos. (D) Immunofluorescence images and corresponding quantification of (E) percentage Ki67+ and (F) SRSF3 negative/low epicardial cells in control and Srsf3 iKO explants, and (G) percentage Ki67+ epicardial cells as a proportion of SRSF3+ and SRSF3 negative/low (SRSF3-/lo) cells in *Srsf3* iKO explants. Scale bar, 50μm. Error bars indicate mean ± SEM (n = 4 - 6). Unpaired t-test with Welch’s correction (*) *P* <0.05, (**) *P* <0.01. (H) PECAM1 whole-mount DAB staining and corresponding quantification (I) of percentage dorsal surface area covered by SV-derived vessels in control and *Srsf3* iKO embryonic mouse hearts at E13.5. Scale bar, 500μm. Error bars indicate mean ± SEM (n = 3). Unpaired t-test with Welch’s correction (*) *P* <0.05. *myo; myocardium, epi; epicardium, LV; left ventricle, endo; endocardium, EPDC; epicardium-derived cells*.

We then examined the major transcriptional changes that occur in *Srsf3*-depleted epicardial cells at E13.5. Whole heart 10X chromium scRNA-seq was performed, as opposed to bulk RNA-seq, since variable recombination was expected in individual epicardial cells^8, 46^. Unsupervised graph-based clustering revealed 18 clusters, largely corresponding to discrete cardiomyocyte types, epicardial, mesenchymal, endocardial, and coronary endothelial cells (Fig. 4A, Supplementary Fig. 3A). The epicardial (Epi) cluster was identified based on mesothelial markers (Supplementary Fig. 3A), such as *Upk3b*^39, 47^. The mesenchymal (Mes) cluster was derived from the epicardium, as indicated by widespread reporter expression (tdTomato; Supplementary Fig. 3B), thus essentially constituting EPDCs. Proportionally fewer mesenchymal cells were recovered in *Srsf3* iKO hearts in comparison to controls (7.4% vs 9.6% of all cells sequenced per genotype; Fig. 4A), consistent with decreased EPDC formation in *Srsf3* iKO hearts (Fig. 3B). In line with primary epicardial cells in explant cultures, FACS-sorted epicardial lineage cells cultured from the same hearts that were processed for scRNA-seq demonstrated a mosaic loss of SRSF3, and reduced proliferation (Cyclin D2 expression) in cells lacking *Srsf3*; Fig. 4B). Since 10X chromium technology has a 3’ bias and detection of gene structural information is limited, we were unable to directly identify absence of exons 2 and 3 in individual cells. However, unexpectedly, upon assessing *Srsf3* levels in the *Srsf3* iKO epicardial population, we detected a bimodal distribution (Fig. 4C). While *Srsf3*-depleted epicardial cells were enriched within the negative to low *Srsf3* expressing population, paradoxically, a substantial proportion displayed *Srsf3* levels that exceeded those in control epicardium (Fig. 4C), noteworthy given the considerable decline in *Srsf3* levels that occurs ordinarily between E11.5-E15.5 (Fig. 1A, B; Supplementary Fig. 1A). Epicardial cells were thus divided into *Srsf3* negative/low or high subsets for downstream analysis. *Srsf3* negative/low epicardial cells showed an overall reduction in predicted proliferation state, with fewer detected in S and G2M phases of the cell cycle in comparison to the *Srsf3* high epicardial cells, based on their expression of cell cycle markers (Fig. 4D). Further, inferred quiescent (G0) or slow-dividing (G1) cells, distinguished by negative *Mki67* expression in the G0/G1 scored population, were enriched in the *Srsf3* negative/low epicardial population (Fig 4E; 90.3% versus 57.9%). Importantly, our analysis inferred that *Srsf3* iKO hearts had a decreased proportion of cycling epicardial cells in comparison to controls (Supplementary Fig. 3C; 9.7% versus 18.8%). *Srsf3*-depleted hearts also demonstrated an increased abundance of epicardial cells with upregulated expression of genes associated with quiescence, such as *Clu*^48^, and senescence, for example *Map4k4*, *Tmem30a* and *Pofut2*^49^ (Fig. 4F). Taken together, these data imply that the *Srsf3*-retaining epicardial cells that evaded recombination would hyper-proliferate and out-compete the *Srsf3*-depleted cells undergoing quiescence or senescence.

**Figure 4.**
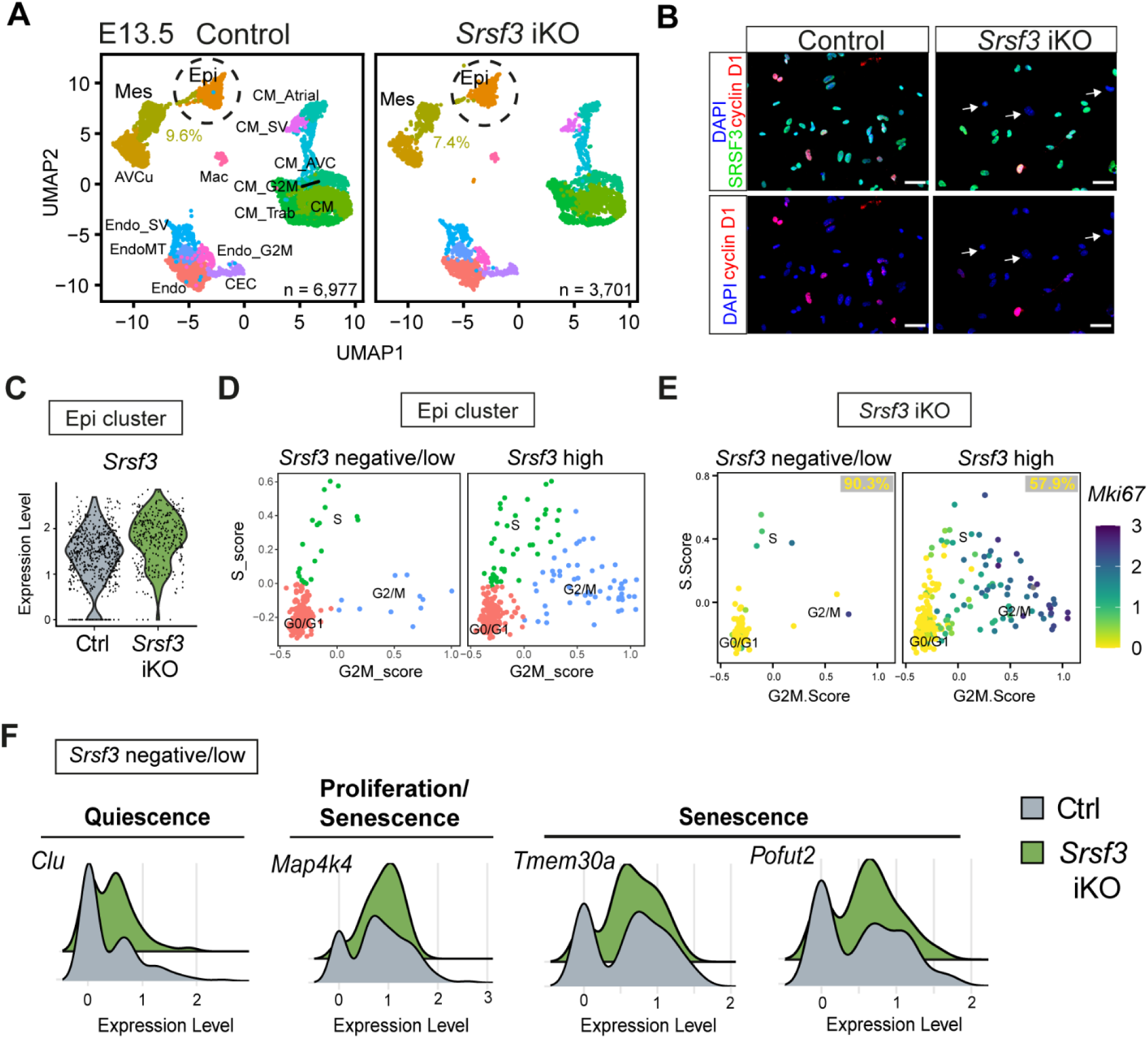
Cell cycle exit and increased senescence of epicardial cells in *Srsf3* iKO hearts. (A) UMAP plots showing the major clusters in control and *Srsf3* iKO embryonic mouse hearts at stage E13.5. Chromium 10X scRNA-seq: control sample; total 6,977 cells, n = 3 hearts, *Srsf3* iKO sample; total 3,701 cells, n = 2 hearts. (B) Immunofluorescence images showing expression of SRSF3 and CCND1 in E13.5 control and *Srsf3* iKO FACS-sorted epicardium lineage (tdTomato+) cells after 48h culture. Scale bar, 50μm. (C) Violin plot showing relative expression of *Srsf3* in the epicardial cluster. (D) Scatter plot showing the distribution of epicardial cells in the indicated cell cycle phases when subsetted by *Srsf3* expression. (E) Feature plot representing range of expression of *Mki67*, further separating quiescent and slow-dividing epicardial cells in *Srsf3* iKO scRNA-seq data at E13.5. Percentage of G0/G1 *Mki67* negative epicardial cells displayed. (F) Histograms representing distribution of *Srsf3*-negative/low expressing epicardial cells for genes related to quiescence (*Clu*), proliferation (*Map4k4*) and sensescence (*Map4k4, Tmem30a* and *Pofut2*) in control and *Srsf3* iKO scRNA-seq data at E13.5. *myo; myocardium, epi; epicardium, LV; left ventricle, LA; left atrium, Endo; endocardial cells, Epi; epicardium, AVCu; atrioventricular cushion, Mes; mesenchymal cells, CM; cardiomyocytes, CM_Trab; trabecular cardiomyocytes, CM_Atrial; atrial cardiomyocytes, CM_AVC; CM atrioventricular canal, Endo_SV; sinus venosus endocardial cells, EndoMT; endocardial-to-mesenchymal transition cells, CEC; coronary endothelial cells, CM_SV; sinus venosus cardiomyocytes, Mac; macrophages,_G2M; proliferating cells*.

Next, we performed differential gene expression analysis guided by one clustering iteration of the Epi cluster in *Srsf3* iKO hearts. This resulted in three subsets: one cluster associated with proliferation genes (Epi_G2M) and two clusters associated with epicardial genes (Epi and Epi2) (Fig. 5A). Notably, the Epi2 subset was enriched for genes associated with hypoxia, such as *Slc2a1* (Fig. 5A, Supplementary Fig. 4A), hypoxia-induced cell death, like *Fam162a*, and attenuated proliferation, such as *Ndrg1* (Fig. 5A, B), an expression signature which was augmented in *Srsf3* iKO epicardial cells. Indeed, the master regulator of hypoxia, *Hif1a*, was found to be more abundant in *Srsf3* negative/low cells (Fig. 5C). Beyond the epicardial phenotype, an upregulation of hypoxia-related *Scl2a1* was detected throughout *Srsf3* iKO hearts (Supplementary Fig. 4A), confirmed by elevated GLUT1 in the compact myocardium (Fig. 5D, E, Supplementary Fig. 4B), possibly as a result of the coronary vessel defects. To further validate the scRNA-seq data, fluorescence *in situ* hybridization (ISH) analysis showed upregulated *Ndrg1* in the epicardium of *Srsf3* iKO hearts (Fig. 5F).^50^. Increased frequency of TUNEL positive apoptotic epicardial cells (Fig. 5G, H) was observed in *Srsf3* iKO hearts compared to controls. The induction of apoptosis in *Srsf3* iKO, but not *Srsf3* cKO epicardium, may reflect the failure of SRSF3-depleted epicardial cells to cope with the hypoxic stress resulting from coronary vessel defects, causing them to undergo cell death. Collectively, these data suggest that SRSF3 regulates epicardial proliferation and survival, to uphold epicardial function through EPDC contribution and promotion of coronary vessel growth.

**Figure 5.**
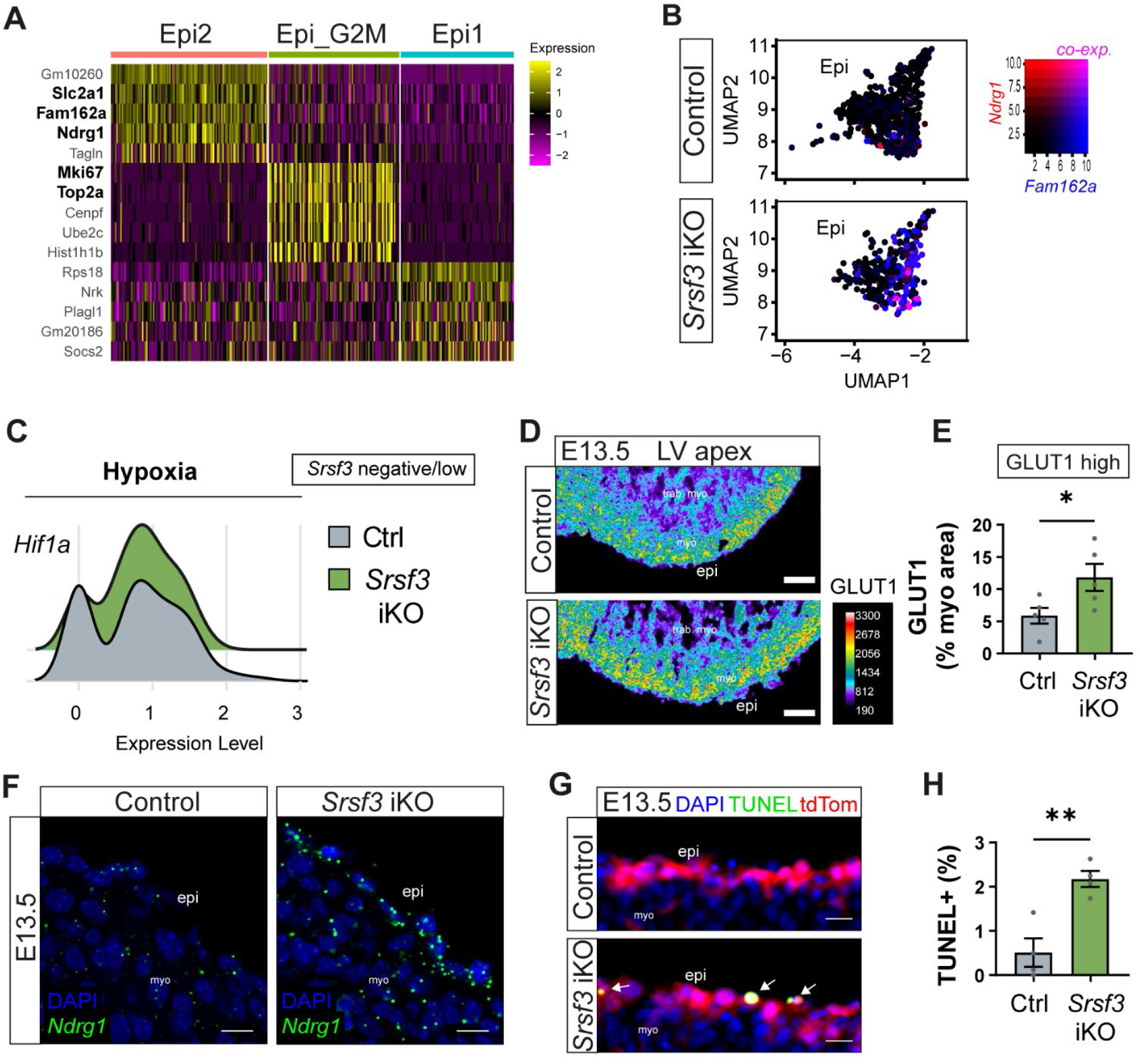
Upregulated hypoxia and increased cell death of epicardial cells in *Srsf3* iKO hearts. (A) Heatmap of top 5 differentially expressed genes for each cluster following one clustering iteration of the Epi cluster in *Srsf3* iKO hearts. High expression is indicated in yellow. Genes of interest highlighted in bold. (B) Feature plots representing range of (co)expression of *Ndrg1* and *Fam162a* mRNA in individual cells of the epicardial (Epi) cluster. (C) Histograms representing distribution of *Srsf3*-negative/low expressing epicardial cells for genes related to hypoxia (*Hif1a*) in control and *Srsf3* iKO scRNA-seq data at E13.5. (D) Pseudocoloured immunofluorescence images and corresponding quantification (E) of GLUT1 high expression as percentage of the ventricular myocardial area in control and *Srsf3* iKO embryonic mouse hearts at E13.5. Scale bar, 75μm. Error bars indicate mean ± SEM (n = 5). Unpaired t-test with Welch’s correction (*) *P* <0.05. (F) Fluorescence ISH of *Ndrg1* mRNA in the epicardium of control and *Srsf3* iKO mouse hearts at E13.5. Scale bar, 10μm. Images representative of n = 4 embryos. (G) TUNEL assay and corresponding quantification (H) of percentage apoptotic (TUNEL+) epicardial cells in control and *Srsf3* iKO embryonic mouse hearts at E13.5. White arrows highlight apoptotic epicardial cells. Scale bar, 20 μm. Error bars indicate mean ± SEM (n = 4). Unpaired t-test with Welch’s correction (**) *P* <0.01. *myo; myocardium, epi; epicardium,trab; trabecular, LV; left ventricle*.

### SRSF3 regulates cell cycle progression in epicardial cells

To further investigate the role of SRSF3 in epicardial cell proliferation and circumvent problems associated with *in vivo* administration of tamoxifen, the active form of tamoxifen, 4-OHT, was used to induce gene deletion *in vitro* in E11.5 epicardial explants (Fig. 6A). Direct treatment resulted in fewer epicardial cells (Fig. 6B) and smaller outgrowth (Fig. 6C) from *Srsf3* iKO explants compared to Wt1^CreERT2^;Rosa26^TdTomato^; *Srsf3*^+/+^ controls. While this strategy enabled more controlled targeting of cells, staining for SRSF3 still revealed mosaic depletion (Fig. 6D). In successfully targeted cells, (SRSF3-/lo), Ki67 expression was diminished (Fig. 6D, E), reinforcing the finding that the proliferative capacity of SRSF3-depleted epicardial cells is impaired.

**Figure 6.**
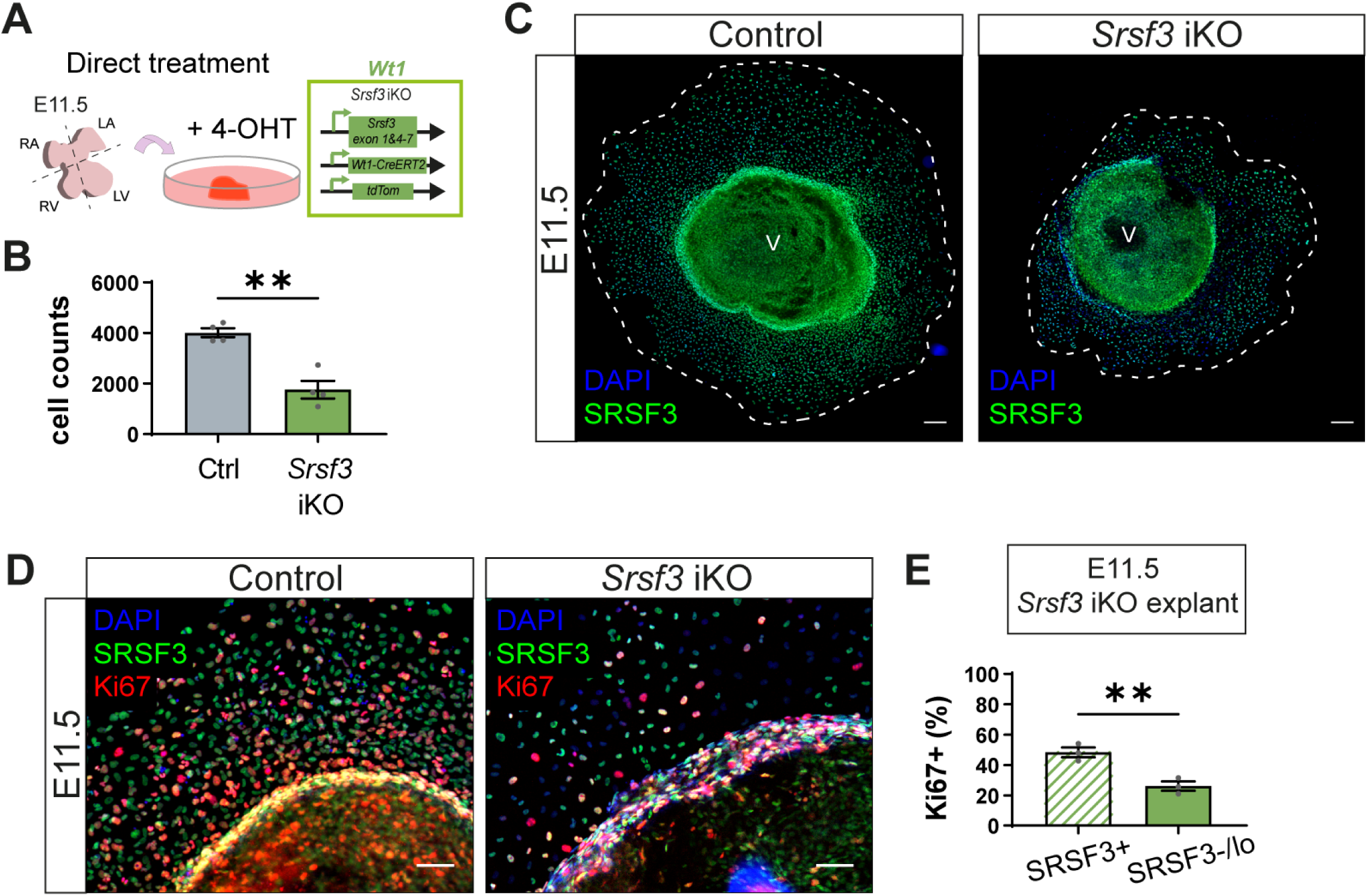
SRSF3 regulates cell cycle progression in primary epicardial cell cultures. (A) Alternate strategy to induce SRSF3 deletion in E11.5 epicardial explants. (B) Quantification of epicardial cells from control and *Srsf3* iKO explants (day 3 culture). Error bars indicate mean ± SEM (n = 4). Unpaired t-test with Welch’s correction (**) P <0.01. (C) Immunofluorescence images showing expression of SRSF3 in control and *Srsf3* iKO explants. Scale bar, 200μm. Image representative of n = 4 embryos. (D) Immunofluorescence images and corresponding quantification of (E) percentage Ki67+ epicardial cells as a proportion of SRSF3+ and SRSF3 negative/low (SRSF3-/lo) cells in *Srsf3* iKO explants. Scale bar, 100μm. Error bars indicate mean ± SEM (n = 3). Unpaired t-test with Welch’s correction (**) *P* <0.01. *LV; left ventricle. RV; right ventricle, V; ventricle, RA; right atrium, LA; left atrium*.

### SRSF3 coordinates epicardial proliferation by controlling key cell cycle regulators and upstream stimulatory pathways

To understand mechanisms through which SRSF3 controls epicardial proliferation, we sought to comprehensively map the SRSF3-RNA interaction networks across the epicardial transcriptome, by infrared CLIP (irCLIP), an ultra-efficient variation of the UV-C crosslinking immunoprecipitation (CLIP) protocol combined with high-throughput sequencing^51^, using a mouse immortalised epicardial cell line^52^ (Fig. 7A; Supplementary Fig. 5A, B). Significant crosslink sites (FDR < 0.05, with short peaks <5nt wide excluded) defined 47,273 peaks of SRSF3 binding mapped to 8,617 different genes across the epicardial transcriptome, with a consensus binding motif matching the one identified in previous SRSF3 iCLIP studies (Fig. 7B)^18, 20, 23^. SRSF3 was found to bind non-coding as well as protein-coding transcripts, with binding sites distributed across predominantly coding sequences and introns, but also regulatory regions (Fig. 7C). Consistent with SRSF3 iCLIP in murine P19 cells^18^, exonic binding was enriched within 100-200nt of 5’ and 3’ splice sites (Supplementary Fig. 5C).

**Figure 7.**
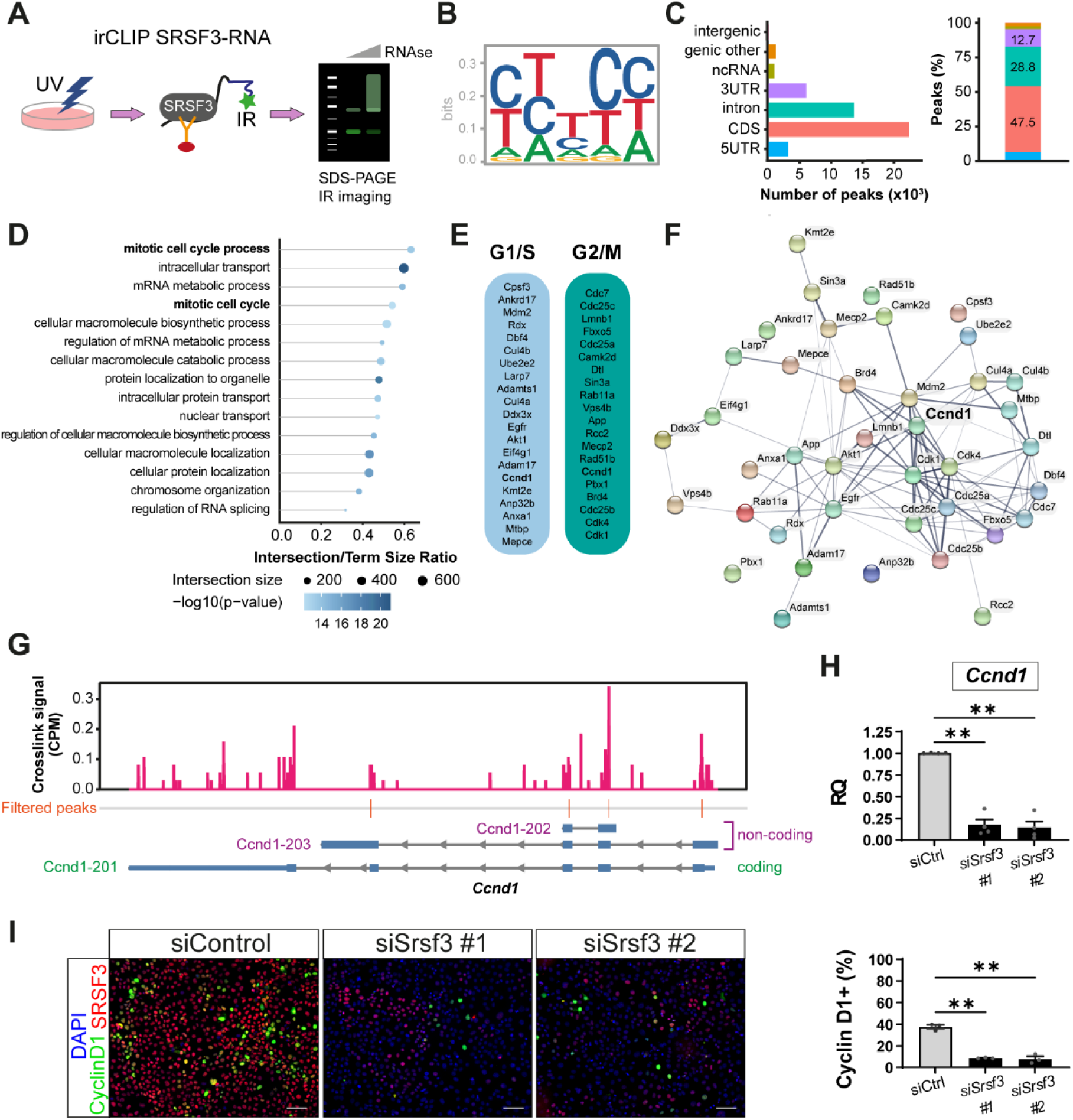
SRSF3 orchestrates proliferation by binding multiple cell cycle regulatory genes and controling *Ccnd1* transcript levels in epicardial cells. (A) Schematic of irCLIP workflow to map direct trancriptome-wide SRSF3-RNA interactions in the epicardial cell line. (B) SRSF3 consensus binding motif in epicardial cells. (C) Number and percentage of significant SRSF3 crosslink peaks (FDR < 0.05) across the transcriptome. (D) GO term analysis on RNA targets identified showing most over-represented biological processes. (E) SRSF3 binding detected to RNAs transcribing ‘positive regulators of G1/S and G2/M transition of mitotic cell cycle’ and (F) corresponding functional association network of these SRSF3 RNA targets in epicardial cells. *Ccnd1*, common in both G1/S and G2/M transition, occupies a central node within the pro-mitotic regulatory network. (G) SRSF3 irCLIP binding profile within *Ccnd1* transcript. Significant peaks, most likely to represent “functional” binding sites were filtered to remove those <5 nt wide (likely transient binding). *Ccnd1* variants are shown (Ccnd1-201, protein coding, Ccnd1-202 and Ccnd1-203, retained intron). (H) qRT-PCR analysis of total *Ccnd1* transcripts in control and *Srsf3* siRNA-transfected cell line; RQ, relative quantification. Error bars indicate mean ± SEM (n = 4). (I) Immunofluorescence images and corresponding quantification of percentage CCND1+ epicardial cells in control and *Srsf3* siRNA-transfected cell line. Scale bar, 100μm. Error bars indicate mean ± SEM (n = 3). One-Way Welch ANOVA and Dunnett’s multiple comparison test (**) *P* <0.01.

Gene Ontology (GO) analysis of genes with significant SRSF3 binding peaks demonstrated an enrichment in functions relating to mitotic cell cycle, RNA metabolism and intracellular transport (Fig. 7D), in keeping with known functions of SRSF3^18, 20, 23^. Among the prominent cell cycle regulators bound by SRSF3 were *Mki67*, *Cdk1*, *Ccnd1* and *Myc* (Supplementary Table 2). Focussing on SRSF3-bound transcripts classified as ‘positive regulators of G1/S and G2/M transition of mitotic cell cycle’ (Supplementary Tables 3 and 4) highlighted canonical regulators, including cyclins, cell division control (Cdc) genes and cyclin-dependent kinases, such as *Ccnd1*, *Cdc7*, *Cdc25a-c*, *Cdk4* and *Cdk1*, as well as upstream inducers of proliferation, such as *Egfr*, *Akt* and *Camk2d* (Fig. 7E). Protein-protein interaction analysis (STRING) of these gene products indicated a functional association network of SRSF3-regulated targets, centred around Cyclin D1 (Fig. 7F). The murine *Ccnd1* gene, comprising five exons, yields a single protein-coding mRNA (*Ccnd1-201*) and the prominent peaks of SRSF3 binding mapped to exons 1, 2, 3 and 4 (Fig. 7G). While splice variants of human *CCND1* are associated with hyper-proliferation and malignancy^53^, there is no evidence for expression of alternatively spliced protein coding *Ccnd1* transcripts in mice^54, 55^. Therefore, the exonic SRSF3 binding identified by irCLIP may rather relate to non-splicing roles in RNA processing, which may influence transcription or stability. Accordingly, we assessed *Ccnd1* expression in epicardial cells after SRSF3 knockdown. Total *Ccnd1* transcript levels were significantly reduced in siSRSF3 transfected cells, compared with controls (Fig. 7H; Supplementary Fig. 5D, E). This corresponded with reduced Cyclin D1 protein expression in knockdown cells (Fig. 7I), and suggests that SRSF3 depletion in epicardial cells induces cell cycle arrest. Collectively, these data support a direct role for SRSF3 in regulating Cyclin D1 expression at the RNA level, over and above any additional degree of control via the upstream regulators identified by irCLIP (Supplementary Table 2), defining SRSF3 as a key orchestrator of epicardial proliferation which acts at multiple levels from signal transduction pathways to the direct effectors of cell cycle progression.

To better understand the mechanisms controlling epicardial cell cycle exit and transition towards senescence, we homed in on SRSF3 targets implicated in cellular senescence (Fig. 4F, Fig. 8A). Specifically, a subset of genes identified by irCLIP are associated with Senescence-Associated Secretory Phenotype (SASP), Oxidative Stress Induced Senescence, DNA Damage/Telomere Stress Induced Senescence, Formation of Senescence-Associated Heterochromatin Foci (SAHF) and Oncogene Induced Senescence (Fig. 8B, Supplementary Tables 5 and 6), via networks in which c-Jun and p53 are central regulators, downstream of MAP kinase signalling (Supplementary Fig. 6A)^56^. While the diversity of targets infers multiple levels of control over cellular senescence by SRSF3, we further investigated *Map4k4*, a 31-exon gene which, in cancer cells, is alternatively spliced by SRSF3, to control c-Jun activation and regulate proliferation^57^. 18 alternatively spliced transcripts of *Map4k4* are reported to exist in human and mouse, via the selective skipping or inclusion of exons 16, 17, 21 and 24, with at least 11 believed to encode proteins (Ensembl). irCLIP revealed SRSF3 binding across the length of the gene (Supplementary Fig. 6B), with notable peaks close to intron-exon boundaries (Fig. 8C). We sought to establish whether alternative splicing of *Map4k4* in epicardial cells is regulated by SRSF3, by knocking down *Srsf3* in the murine MEC1 epicardial cell line^58^. Whilst total levels of *Map4k4* were unaffected (Fig. 8D), *Srsf3* depletion resulted in significant alternative splicing, in favour of an overall increase in the skipping of exon 17, a key exon implicated from cancer studies in the control of proliferation (Fig. 8E, F). Specifically, in the absence of *Srsf3*, exon 16 skipped:exon 17 retained isoforms (*Map4k4-206, 208, 209*) were reduced, whilst exon 16 retained:exon 17 skipped isoforms (*Map4k4-202, 215, 217*) were increased, as were isoforms lacking both exons 16 and 17 (*Map4k4-205, 213*). Isoforms with both exons retained (*Map4k4-201, 218*) were lower in abundance in MEC1 cells at baseline, and unchanged by *Srsf3* knockdown. Taken together, these data demonstrate a role for SRSF3 in the alternative splicing of *Map4k4*, to alter the balance between isoforms that are known to preferentially promote proliferation^59^ or senescence^60^, suggesting that regulation by SRSF3 may, at least in part, drive the switch of epicardial cells from the proliferative to senescent state.

**Figure 8.**
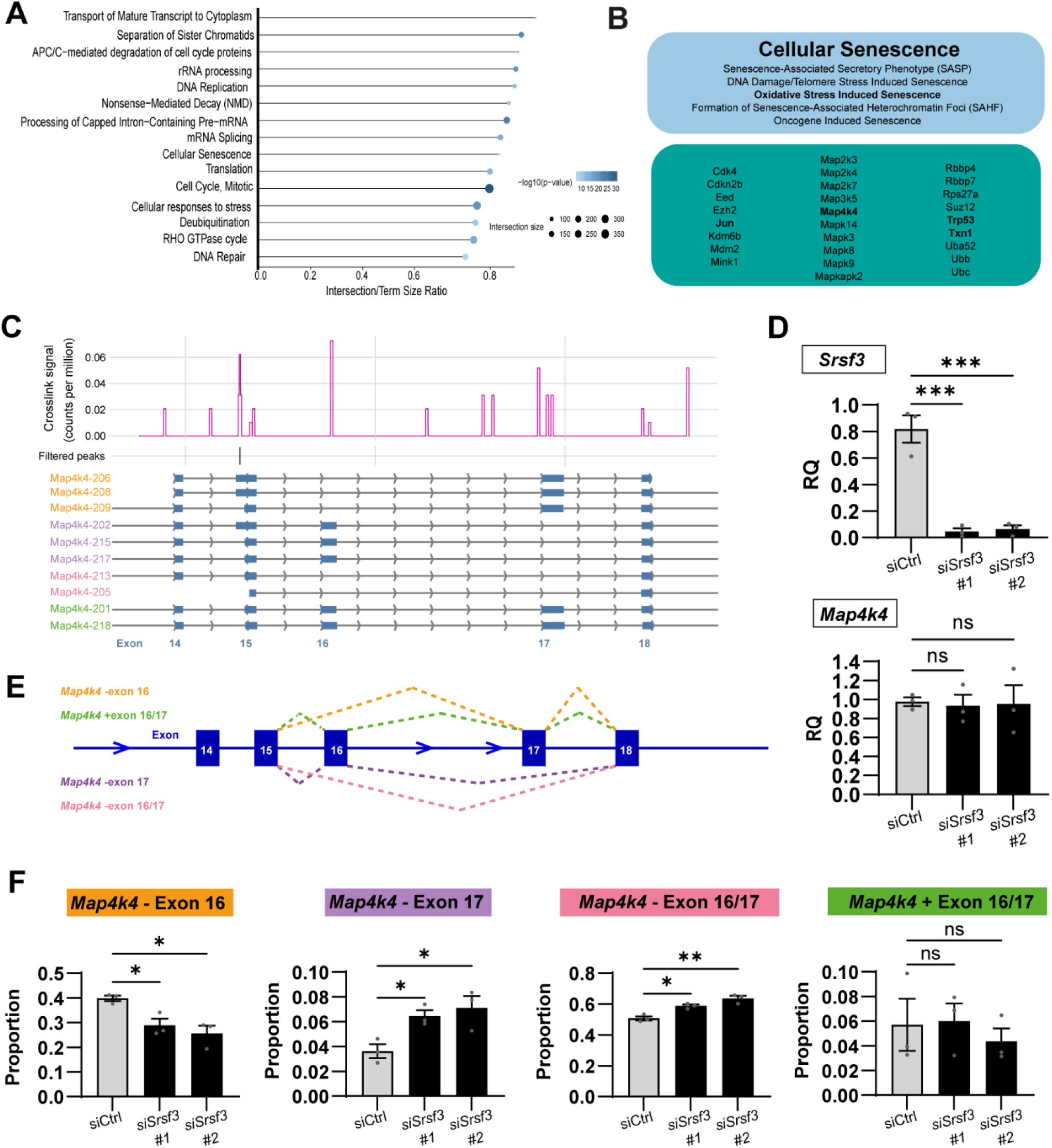
SRSF3 controls the alternative splicing of *Map4k4*, a gene that regulates cell proliferation and senescence. (A) Reactome analysis on RNA targets identified by irCLIP, showing highly represented biological processes. (B) SRSF3 target genes associated with cellular senescence (sub)terms. (C) SRSF3 irCLIP binding profile across exons 14-18 of the *Map4k4* gene), to highlight SRSF3 binding sites around key exons 16 and 17, and the protein coding isoforms subject to alternative splicing in this region. (D) qRT-PCR analysis of *Srsf3* and total *Map4k4* transcripts in control and *Srsf3* siRNA-transfected MEC1 cell line; RQ, relative quantification. (E) Schematic of splicing events assessed by qRT-PCR after *Srsf3* knockdown. (F) qRT-PCR analysis of *Map4k4* splicing in control and *Srsf3* siRNA-transfected MEC1 cell line; RQ, relative quantification. Error bars indicate mean ± SEM (n = 3 separate experiments). One-Way Welch ANOVA and Dunnett’s multiple comparison test (*) *P* <0.05; (**) *P* <0.01.

### Non-recombined epicardial lineage cells compensate for loss of SRSF3-depleted cells

Having identified defects in epicardial proliferation and viability in Srsf3-depleted cells, contrasting with the retention or up-regulation of Srsf3 levels in non-recombined cells at E13.5, we sought to determine the impact of mosaic targeting on later stage heart development. We first assessed the tdTomato positive epicardial lineage of *Srsf3* iKO and control hearts at E15.5 using SMART-Seq2^61^. Somewhat surprisingly, by this stage, the *Srsf3* iKO lineage displayed similar composition to controls, consisting of a small epicardial cluster, two mesenchymal clusters, and a distinct cluster of mural cells (Fig. 9A, Supplementary Fig. 7A). We previously characterised Mes1 as subepicardial mesenchyme and Mes2 as fibroblast-like cells^39^. Since the plate-based SMART-Seq2 method allows recovery of full-length mRNA, we could establish that no recombined cells lacking *Srsf3* exons 2/3 were present at this stage, indicating that *Srsf3*-depleted cells must have been eliminated earlier in development. Consistent with previous reports^8, 46, 62^, it should be noted that tdTomato expression does not accurately correlate with gene depletion, due to the more efficient recombination of the shorter transcriptional stop sequence of Cre reporters compared with recombination across multiple exons within floxed alleles. Therefore, to confirm that only non-*Srsf3*-recombined cells remained, E15.5 explants were stained for SRSF3 (Supplementary Fig. 7B). In contrast to E13.5 explants, all outgrowing epicardial cells at E15.5 had high levels of SRSF3, confirming that the SRSF3-depleted cells had been removed earlier in development. In fact, the SMART-Seq2 data indicated increased *Srsf3* expression in both epicardial and subepicardial mesenchymal cells of *Srsf3* iKO hearts, compared with controls, which was confirmed by SRSF3 IHC (Fig. 9B, C, Supplementary Fig. 7C). Thus, the modest elevation of Srsf3 in non-recombined epicardial cells initially detected at E13.5 is robustly maintained, and even enhanced, across a developmental window through to at least E15.5. The increased SRSF3 expression coincided with increased proliferative capacity, as indicated by *Top2a* expression and confirmed by TOP2α staining in the epicardium (Fig. 9D, E, Supplementary Fig. 7D). We found that the reduced incidence of EPDCs invading the myocardium from E13.5 through to E15.5 (Fig. 3B, 9F, Supplementary Fig. 7E) was fully restored by E17.5 (Fig. 9G, Supplementary Fig. 7E), and the earlier compaction defect rescued, to leave hearts appearing morphologically normal by this late stage in development (Supplementary Fig. 7F). Together, these data indicate that the epicardial cells that evade recombination possess a remarkable capacity to compensate for the early loss of SRSF3-depleted cells, by using SRSF3-controlled mechanisms to promote epicardial proliferation and restore normal function.

**Figure 9.**
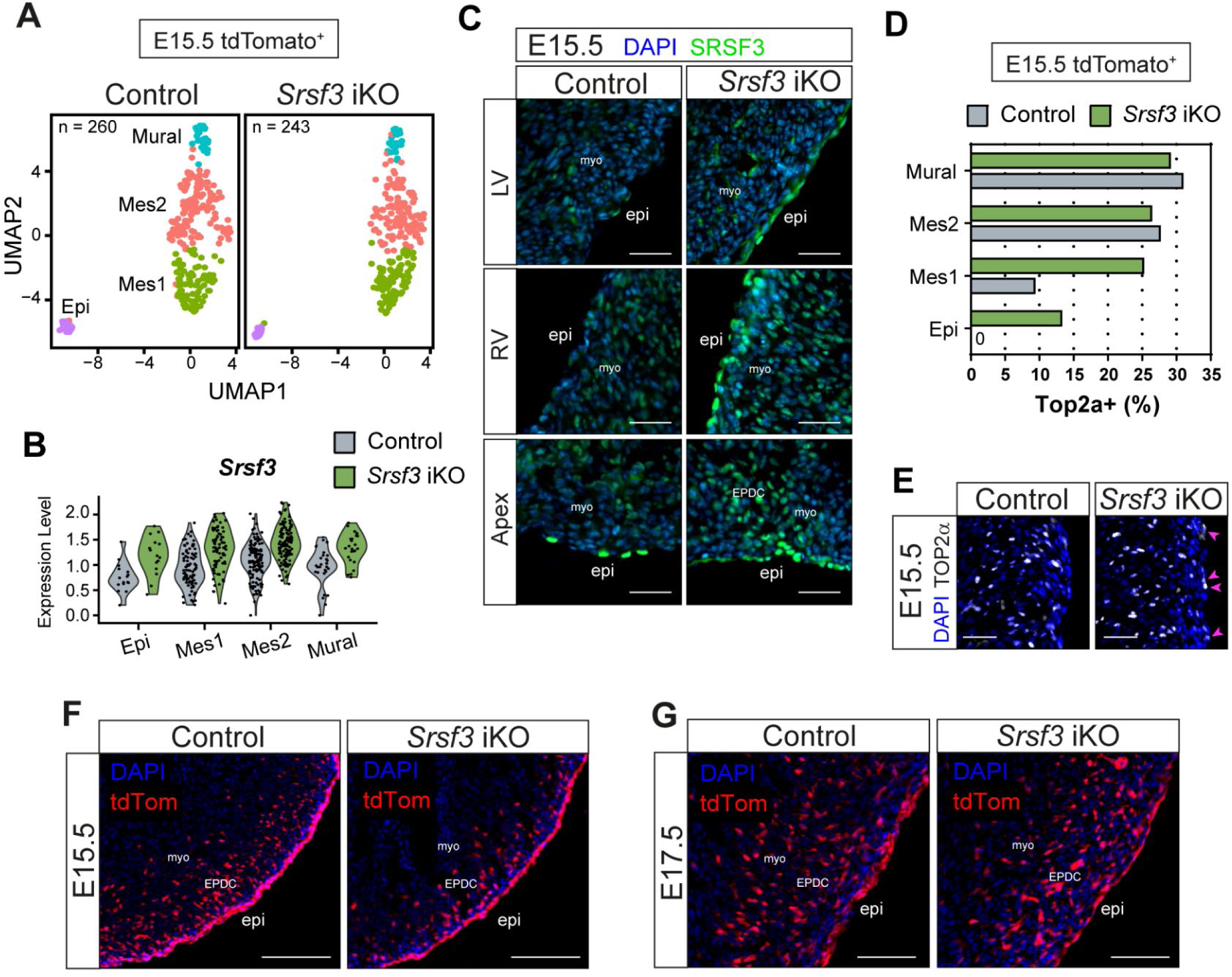
Non-recombined cells of the epicardial lineage upregulate SRSF3 and hyperproliferate to compensate for loss of SRSF3-depleted cells in *Srsf3* iKO hearts. (A) UMAP plots showing the major cell clusters in the epicardial lineage of control and *Srsf3* iKO embryonic mouse hearts at E15.5. FACS-sorted epicardium lineage (tdTomato+) cells plate-based scRNA-seq: control sample; total 260 cells, n = 6 hearts, *Srsf3* iKO sample; total 243 cells, n = 4 hearts. (B) Violin plot showing relative expression of *Srsf3* in individual clusters. (C) Immunofluorescence images showing expression of SRSF3 in the epicardium of control and *Srsf3* iKO embryonic mouse hearts at E15.5. Scale bar, 50μm. Images representative of n = 4. (D) Quantification of percentage *Top2a*-expressing epicardial lineage cells in individual clusters of control and *Srsf3* iKO scRNA-seq data at E15.5. (E) Immunofluorescence images showing expression of TOP2α in the epicardium of control and *Srsf3* iKO embryonic mouse hearts at E15.5. Magenta arrowheads highlight upregulated expression of TOP2α in epicardial cells. Scale bar, 50μm. Images representative of n = 3. (F, G) Fluorescence images showing epicardial lineage cells (tdTomato+) in control and *Srsf3* iKO embryonic mouse hearts at E15.5 (F) and E17.5 (G). Scale bar, 100μm. Images representative of n = 3 embryos. *myo; myocardium, epi; epicardium, EPDC; epicardium-derived cells, Epi; epicardial cells, Mes; mesenchymal cells*.

### SRSF3 depletion in epicardial progenitors leads to compaction and coronary vasculature defects

To explore the extent to which compensatory mechanisms could overcome more severe early epicardial deficiencies, we tested two alternative strategies to achieve more efficient gene deletion in the *Srsf3* iKO model with a higher dose of tamoxifen (80mg/kg): at i) E8.5, expected to target progenitors in the PEO; ii) E9.5, expected to improve targeting of epicardial founders. We observed that induction at E8.5 resulted in a more severe myocardial non-compaction in some hearts, as revealed by the persistence of a highly trabeculated EMCN+ endocardium, which extended closer to the epicardial surface (Fig. 10A; Supplementary Fig. 8A). However, given the variable extent of compensation and limited accuracy of histological morphometric analysis, this phenotype was not statistically significant overall (Fig. 10B). However, the vascular defects remained at E17.5 after E8.5 induction, evidenced by an increased proportion of superficial vessels, relative to those that had invaded the myocardium, in *Srsf3* iKO hearts compared to controls (quantified as % lumenised vessels within the 50μm of myocardium underlying the epicardium; Fig. 10A, C; Supplementary Fig. 8A; magenta arrows). Impaired coronary vasculature formation, notably the appearance of ectopic subepicardial vessels, frequently accompanies myocardial non-compaction in hearts with compromised epicardial function^63–65^. Although epicardial formation and function is initially impaired with all tamoxifen regimes tested, each showing disrupted morphology, with rounded cells on the surface and limited invasion, the respective E17.5 phenotypes appear to reflect the differential capacity for compensation, which is greater in the E9.5/E11.5 regime, compared with E8.5. Efficient compensation and phenotypic rescue was similarly observed in *Srsf3* iKO hearts with 80mg/kg tamoxifen at E9.5, comparable to the E9.5/E11.5 phenotype (Supplementary Fig. 8A), suggesting that timing of administration determines the extent of *Srsf3* recombination and functional impairment. Taken together, these data indicate that the transient functional impairments observed with the original tamoxifen regime persist when a higher efficiency SRSF3 depletion is achieved (Fig. 10, Supplementary Fig. 8A), due to the essential role of cell division in epicardial function^16^.

**Figure 10.**
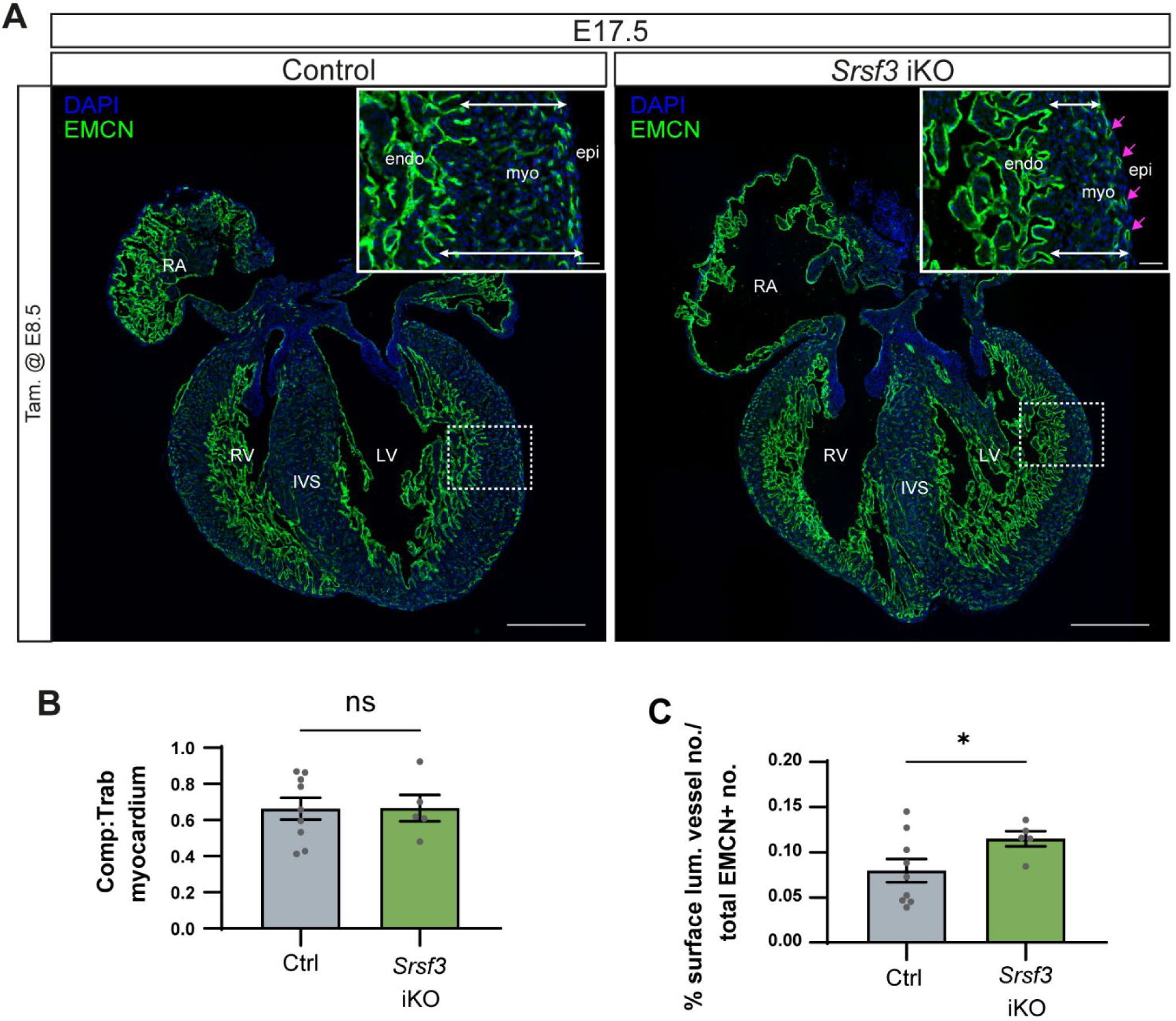
Defects in coronary vasculature persist with early-stage induction of SRSF3 depletion in Srsf3 iKO hearts. (A) Immunofluorescence of the endocardium and vascular network (EMCN+) in control and *Srsf3* iKO embryonic mouse hearts at E17.5. Tamoxifen induction was administered at E8.5. Inset shows selected region of interest of the left ventricular wall. White arrows highlight myocardial wall thickness and magenta arrows highlight presence of superficial vessels. Scale bar, 500μm and inset 50μm. Images representative of n = 5-9 embryos. (B) Myocardial compaction, quantified as compact:trabeculated myocardial ratio. (C) Percentage surface lumenised vessels within 50 µM of the epicardium, as a % of total number of lumenised vessels. Error bars indicate mean ± SEM (n = 5-9 embryos). Unpaired t-test with Welch’s correction (*) *P* <0.05. *LV; left ventricle, RV; right ventricle, RA; right atrium, IVS; intraventricular septum, myo; myocardium, epi; epicardium, endo; endocardium*.

## Discussion

Our study presents insight into the critical role of SRSF3 in mediating proliferation of the (pro)epicardium, to enable proper heart development. We show that SRSF3 expression is widespread in embryonic hearts, with highest levels detected in proepicardial cells and in the epicardial layer until E12.5. Constitutive SRSF3 depletion in epicardial progenitor cells resulted in gross morphological heart abnormalities and embryonic lethality at E12.5. Failure to form the epicardial layer was due to a diminished source of epicardial progenitor cells within the PEO, as a result of their decreased proliferative capacity in the absence of SRSF3. However, recombination in the Tg(Gata5-Cre) model also targeted a subset of cardiomyocytes. Thus, combined depletion of SRSF3 in most epicardial progenitor cells and a subset of cardiomyocytes contributed to the severe phenotype, since embryos with compromised epicardial formation generally die at later stages^66, 67^. Consistent with our findings, early stage embryonic lethality was previously reported in cardiac-specific SRSF3 knockout mice due to impaired cardiomyocyte proliferation and survival^34^.

Notably, peak epicardial SRSF3 expression at E11.5 coincides with the reported peak of epicardial proliferative activity^16^. Temporal Cre-induction and selective deletion of *Srsf3* in epicardial cells, using the Wt1^CreERT2^ line, resulted in impaired proliferation, ultimately leading to loss of SRSF3-depleted epicardial cells from the heart. SRSF3 was key to epicardial cell cycle progression as cyclin D1, found to be an SRSF3 target gene, was downregulated in SRSF3-depleted cells *in vitro*. Beyond regulation of a key cyclin, our irCLIP experiments mapped a significant proportion of SRSF3-RNA binding sites to a range of proliferation-associated transcripts in epicardial cells. To our knowledge, we present the first endogenous SRSF3 CLIP analysis, identifying RNA binding targets without the confounding influence of protein tagging and over-expression. Importantly, this analysis validated SRSF3 as a positive regulator of G1 to S phase transition in epicardium, consistent with roles demonstrated in other cell types^28, 30^. A salient finding from our study was that the epicardial cells which escaped recombination and hyper-proliferated to restore epicardial function, were those with elevated SRSF3 levels, compared with control, reinforcing the notion that SRSF3 critically drives cell cycle in the epicardium.

Of note, the role of SRSF3 in driving cancer progression has been investigated in the context of *MAPK4K* alternative splicing^57, 68^. In the most aggressive cancers, with the highest levels of *SRSF3,* high proliferation rates correlated with prominent expression of exon 16/17-containing *MAP4K4* isoforms; in contrast, cancers expressing lower levels of *SRSF3* proliferated more slowly and associated with *MAP4K4* isoforms in which these exons were predominantly skipped^57^. *SRSF3* was directly implicated in the proliferation and invasiveness of U2OS and HeLa cancer cells, with *SRSF3* knockdown inhibiting cell proliferation, associated with *MAP4K4* exon skipping^68^. We now demonstrate that SRSF3 engages similar mechanisms of alternative splicing to drive cellular proliferation during embryonic development, as it does in oncogenesis.

SRSF3 depletion in the epicardium resulted in cell death associated with hypoxic stress, exemplified by upregulated *Fam162a* and an increase in TUNEL+ epicardial cells. *Fam162a* is a transcriptional target of HIF-1α, which promotes its expression under hypoxic conditions^69^.. Although we detected SRSF3 binding to *Fam162a* by irCLIP, we cannot distinguish the extent of direct regulation versus indirect mechanisms, which may be induced as a secondary consequence of hypoxic stress in *Srsf3* iKO hearts, due to vascular insufficiency as a result of inadequate epicardial stimulation of sinus venosus sprouting. It may be of relevance that acute hypoxia in tilapia fish resulted in alternative splicing of *Fam162a*^70^ and that de-regulated expression of RNA splicing factors such as SRSF3, were reported to mediate numerous alternative splicing events to promote cancer cell evasion of apoptosis^71^. Either way, SRSF3 depletion may expose epicardial cells to hypoxia-induced apoptosis, and the relationship between SRSF3 and *Fam162a* warrants further investigation.

Unlike targeting with Tg(Gata5-Cre), induction of SRSF3 deletion using Wt1^CreERT2^ at E8.5, did not prevent epicardial formation, although it delayed formation. This could be due either to the reduced efficiency of the inducible Cre or the timing of Cre activation, which occurs at the onset of proepicardial formation with Tg(Gata5-Cre), compared with Wt1^CreERT2^, which suffers from a lag in recombination after tamoxifen delivery^72^. Although the epicardial layer was established in *Srsf3* iKO hearts, epicardial function was clearly defective, manifested as failed EMT, myocardial non-compaction and impaired coronary vessel formation, consistent with other studies^63, 64, 67, 73^. In contrast, induction of SRSF3 depletion at E9.5 or E9.5/ E11.5 resulted in normal heart morphology at E17.5, due to the ability of non-recombined cells to over-express SRSF3 and hyperproliferate to restore the epicardial lineage. The compensatory mechanism is probably facilitated by the loss of SRSF3-depleted cells coupled with cell competition mechanisms that are active in the epicardium, as previously demonstrated with overexpression of c-MYC^74^. c-MYC overexpressing cells similarly become hyperproliferative and out-compete wild-type cells in epicardial explants^74^. Alternatively, hypoxic stress generated by diminished SV sprouting may upregulate SRSF3 in non-recombined cells, in line with reported hypoxia-induced SRSF3 expression^75^, to enhance their proliferation. Thus, this study reveals an extraordinary potential of epicardial cells to hyperproliferate and compensate in the presence of genetic change. It also highlights the limitations of variable recombination efficiency, which may confound investigation of critical genes in the maturing epicardium. A significant number of studies use inducible Cre lines without considering the consequences of mosaic targeting, and accordingly risk misinterpreting data, especially when more readily recombined reporters such as tdTomato are used as surrogates for target gene recombination. The impact of incomplete recombination is compounded in the Wt1CreERT2 lines, given the narrow developmental window for specific induction^39^ and the potent cell competition mechanisms that drive enrichment of non-targeted cells during development. We highlight the shortcomings and key considerations exposed in our study, to promote the appropriate use of inducible genetic models, and the rigour and reliability of future investigations.

Our study adds to current literature describing the key role of SRSF3 in cellular proliferation^27, 28, 30, 37, 68, 76, 77^. In the context of cardiac development, SRSF3 is required for the proliferation and survival of (pro)epicardial cells. SRSF3 regulation of cell cycle progression is essential to enable epicardial cells to undergo EMT and to subsequently support vascular development. This study defines SRSF3 as a key regulator of epicardial cell behaviour during development. Moreover, the finding that compensatory restoration of the epicardium can be achieved by induction of cell cycling and the delineation of the molecular mechanisms controlling epicardial proliferation may provide relevant insights for regenerative therapies.

## Materials and methods

### Mouse strains

Conditional and inducible targeting of SRSF3 was achieved by crossing epicardial Cre lines. Tg(Gata5-Cre)1Krc^40^ and Wt1^tm2(cre/ERT2)Wtp 43^, respectively, with mice in which exons 2 and 3 of *Srsf3* were flanked by loxP sites^34, 35^. The epicardial lineage was traced using Rosa26tdTomato in crosses with Gt(ROSA)26Sor^tm14(CAG-tdTomato)Hze 41^. Controls used were either Cre+; Srsf3+/+ or littermate Srsf3fl/+ (tamoxifen induced, in the case of iKO), or Srsf3fl/fl; Cre-. Pregnant females were oral gavaged with 40mg/kg at E9.5 and E11.5, or 80mg/kg tamoxifen at E8.5 or E9.5 (where specifically stated). All procedures were approved by the University of Oxford Animal Welfare and Ethical Review Board, in accordance with Animals (Scientific Procedures) Act 1986 (Home Office, UK).

### Tissue harvest and cryosectioning

Embryos and embryonic hearts were harvested and fixed in 4% PFA for 2h at room temperature (RT). Following PBS washes, tissues were equilibrated overnight at 4°C in 30% sucrose/PBS, and then gradually transitioned into O.C.T. Samples were stored at −80°C, then cryosectioned at 8-12µm thickness.

### (Pro)epicardial explant and cell line culture

Embryonic hearts were dissected into atrial and ventricular pieces and cultured on 1% gelatine-coated dishes in culture medium (15% FBS, 1% Penicillin/Streptomycin, DMEM high glucose Glutamax). Explants were incubated at 37°C in 5% CO_2_ for 3 days. Dissected proepicardia were subjected to the same culture conditions as above. The strategy to induce SRSF3 depletion *in vitro* involved culturing epicardial explants in culture medium supplemented with 1µM 4-Hydroxytamoxifen (4-OHT). An immortalised epicardium-derived cell line^52^ was cultured on 1% gelatine in immorto medium (10% FBS, 1% Penicillin/Streptomycin, ITS supplement, 0.1% IFNγ, DMEM high glucose Glutamax) at 33°C in 5% CO_2_. The MEC1 epicardial cell line^58^ was cultured on 0.1% gelatine in DMEM high glucose with Glutamax, 10% FBS, 1% Penicillin/Streptomycin, at 37°C in 5% CO_2_. Cells were transfected with 40nM or 80nM siRNA (siControl; silencer siRNA negative control Cat. #4390843, siSrsf3 #1; s73613, siSrsf3 #2; s73615, ThermoFisher) for 48h using Lipofectamine RNAiMAX Transfection Procedure (ThermoFisher Cat.#13778-100).

### Immunofluorescence

Standard protocols were used. Full details can be found in Supplementary Methods.

### RNA extraction and qRT-PCR

Full details can be found in Supplementary Methods.

### Flow cytometry

Full details can be found in Supplementary Methods.

### Single-cell RNA-sequencing and analysis

A full description of the experimental and analysis methods can be found in Supplemental Methods.

### Epicardial Srsf3 irCLIP and Analysis

irCLIP was performed using the epicardial cell line, largely as described in^51, 78–82^ with some adaptations of the protocol. Primers were ordered as published from IDT. The L3-biotin pre-adenylated linker was ordered with the IRdye-800CW attached from IDT: /5rApp/AGATCGGAAGAGCGGTTCAGAAAAAAAAAAAA/iIRD800CWN/AAAAAAAAAAAA/3Bio/. Cells were washed with PBS and crosslinked with 200mJ/cm2 UV light in a Stratalinker 1800. Protein A Dynabeads (Life Technologies) were washed and coated with 10ug anti-SRSF3 antibody (ab198291, Abcam). Cells were resuspended in lysis buffer (50 mM Tris-HCl, pH 7.4, 100 mM NaCl, 1 % Igepal CA-630, 0.1 % SDS, 0.5 % sodium deoxycholate, protease inhibitor (Sigma Aldrich)) and samples sonicated in a Bioruptor (Diagenode). In the presence of Turbo DNase (Thermo Scientific), RNA in 100ug lysate per sample was partially digested with 1/500 RNase I or 1/20 RNaseI (Invitrogen) for the high RNase I control sample. SRSF3 and crosslinked RNA were immunoprecipitated from lysate for 2h at 4°C with rotation. Beads were washed once each with high stringency buffer (20 mM Tris, pH 7.5, 120 mM NaCl, 25 mM KCl, 5 mM EDTA, 1% Trition-X100, 1% Na-deoxycholate), high salt buffer (20 mM Tris, pH 7.5, 1 M NaCl, 5 mM EDTA, 1% Trition-X100, 1% Na-deoxycholate, 0.001% SDS) and low salt buffer (20 mM Tris, pH 7.5, 5 mM EDTA). RNA was dephosphorylated with polynucleotide kinase (NEB) at 37°C for 30min, followed by washes with high salt buffer and NT2 buffer (50 mM Tris, pH 7.5, 150 mM NaCl, 1 mM MgCl_2_, 0.0005% Igepal). The L3-App-IRD800CWN-biotin linker was ligated to the RNA with T4 RNA ligase (NEB) overnight at 16°C. Beads were washed with high salt and NT2 buffer and eluted in NuPAGE loading buffer (Invitrogen) and run on a NuPAGE 4-12% Bis-Tris gel (Invitrogen) and transferred onto a 40μm nitrocellulose membrane. The SRSF3-RNA complexes were visualised with an Odyssey Clx infrared imager. Protein-RNA complexes containing RNA of approximately 70-280nt length (20-80kDa above the protein-RNA-adapter complex) were excised and RNA released by proteinase K (Roche) digestion for 60min at 50°C in proteinase K digestion buffer (100 mM Tris, pH 7.5, 50 mM NaCl, 1 mM EDTA, 0.2% SDS). RNA was extracted with phenol-chloroform (Sigma Aldrich) and precipitated with ethanol overnight. RNA was reverse transcribed with TIGRT-III reverse transcriptase (Ingex) and cDNA/RNA hybrids captured with MyOne Streptavidin C1 Dynabeads (Life Technologies). RNA was degraded by alkaline hydrolysis for 15min in RNA degradation buffer (0.1 μM P3short oligonucleotide, 3.75 mM MnCl_2_) followed by cDNA circularization with Circligase II (Epicentre). Circularized cDNA was captured on Ampure XP beads (Beckman Coulter) before elution in water. The cDNA was amplified with Phusion HF polymerase (NEB) and P3/P6 primers in a ViiA7 qPCR machine in the presence of SYBR Green (Thermo Scientific) for real-time visualisation of the amplification product. The library was cleaned-up with Select-a-Size DNA Clean & Concentrator kit (ZymoResearch) to remove empty library before a further three PCR cycles using P3/P6Solexa primers to add the adapters necessary for Illumina sequencing. The final library was cleaned up first with Ampure XP beads followed by gel clean-up on a Novex TBE-6% Urea gel. Libraries were visualised with SYBR Gold (Thermo Scientific) and excised to remove residual empty library. DNA was extracted from the gel in Crush-Soak Gel buffer (500 mM NaCl, 1 mM EDTA, 10 mM Tris pH 7.5, 0.1% SDS) overnight at 55°C and cleaned up using DNA Clean-and-Concentrator-5 kit (Zymo Research). Libraries were quantified with KAPA Library Quantification Kit (Roche) and sequenced on a NextSeq 550 with a High Output Kit v2 (75 Cycles) (Illumina).

Sequencing data were demultiplexed with Cutadapt 2.1 (Martin, 2011, EMBnet.journal) and processed with the nf-core/clipseq pipeline^79^. Crosslinks from the three replicates were merged and used as input to iCount-Mini^82^ using an FDR threshold of < 0.05 as significant. Peaks less than 5 nt wide were filtered out. For motif analysis, we examined the top 20 pentamers occurring in the −20…+20 window around the peak starts. GO term enrichment analysis was performed using gprofiler2 (v0.2.3)^78^ using genes with >10 peak crosslinks and a background set defined as all genes with crosslinks. Reactome analysis was done using ReactomePA package (v1.46.0)^83^. IGV viewer^82^ was utilised to browse irCLIP data and clipplotr ^84^ was utilised for irCLIP data visualisation.

### Whole-mount DAB staining

Full details can be found in Supplementary Methods.

### Western Blot

Full details can be found in Supplementary Methods.

### Fluorescence in situ hybridization (mRNA)

RNAscope Multiplex Fluorescent v2 assay (ACD) was performed on cryosections according to manufacturer’s instructions, with minor modifications as previously described ^39^. Ndrg1 and negative control probe, and TSA plus fluorophores, are listed in Supplementary Methods Table 3.

### TUNEL assay

Click-iT™ Plus TUNEL Assay (ThermoFisher) was performed on cryosections according to the manufacturer’s instructions.

### Data access

All sequencing data from this study are available in Gene Expression Omnibus database under accession number GSE145832, GSE205797 and E-MTAB-11853.

### Statistical analysis

Statistical analysis was performed in GraphPad Prism 9 software. Unpaired t-test with Welch’s correction was used to compare two experimental groups. One-way Welch ANOVA and Dunnett’s multiple comparison test was used to compare more than two experimental groups. P value lower than 0.05 was considered significant. Details for statistical analyses, including biological replicate numbers, are included in figure legends.

## Acknowledgements

We thank Prof Peter Nielsen, Max Planck Institute, and Prof Enrique Lara-Pezzi, CNIC, for the Srsf3^fl/fl^ line; Prof William Pu, Harvard, for the Wt1^CreERT2^ line; Prof Kenneth Chien, Karolinska Institutet, for the Tg(Gata5-Cre) line, Dr Madeleine Lemieux (Bioinfo) for processing SMART-Seq2 and 10X data; Dr Neil Ashley, WIMM Single cell facility, for sequencing, the Dunn School Flow Cytometry facility for equipment use, Micron for microscopy facilities and Biomedical Services staff for animal husbandry. This work was funded by the British Heart Foundation (BHF): DPhil Studentship (FS/15/68/32042); BHF Project grant (PG/15/112/31940); BHF Ian Fleming Senior Basic Science Research Fellowships (FS/13/4/30045 and FS/19/32/34376); BHF Centre of Regenerative Medicine, Oxbridge (RM/13/3/30159).

## Author contributions

Conceptualisation, I-EL, NS; Methodology, I-EL, SB, AMC, IRM, TC, ANR, NS; Investigation, I-EL, ANR, SB, NS; Data Analysis, I-EL, SB, AMC, IRM, TC, ANR, NS; Writing – Original Draft and Revisions, I-EL, ANR, NS; Writing – Review and Editing, all authors; Funding Acquisition, NS; Supervision, ANR and NS.

