## Supplementary Information for "SRSF3 is a key regulator of epicardial formation"

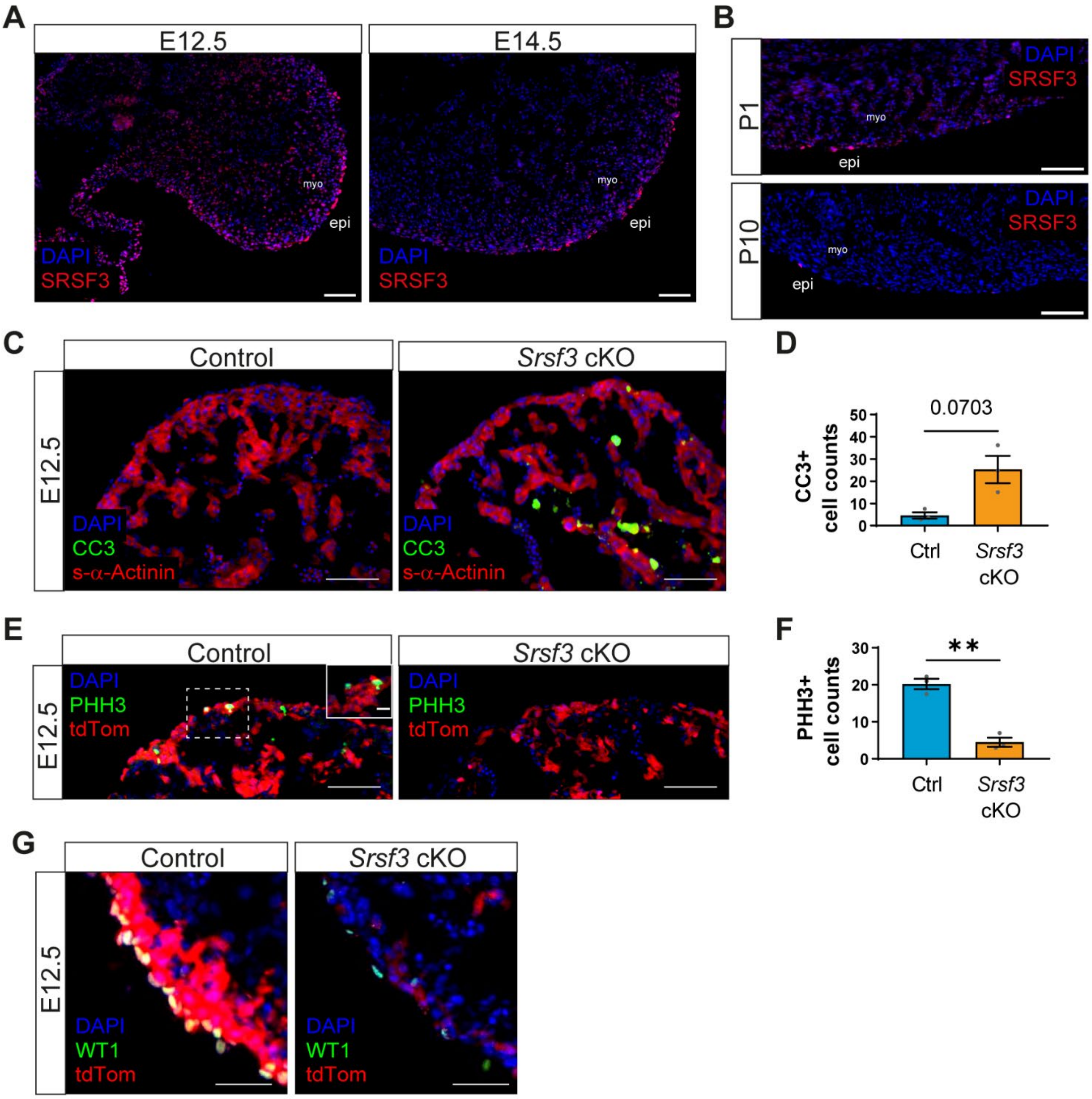

**Supplementary Figure 1. Increased cell death and decreased proliferation in *Srsf3* cKO hearts.** (A) Immunofluorescence cryosection images showing expression of SRSF3 in embryonic mouse hearts at E12.5 and E14.5. Scale bar, 100µm. Images representative of n = 4 embryos. (B) Immunofluorescence cryosection images showing expression of SRSF3 in postnatal mouse hearts at P1 and P10. Scale bar, 100µm. Images representative of n = 4 hearts. (C) Immunofluorescence images and corresponding quantification (D) of CC3+ cells in control and *Srsf3* cKO embryonic hearts at E12.5. Scale bar, 50µm. Error bars indicate mean ± SEM (n = 3). Unpaired t-test with Welch's correction. (E) Immunofluorescence images and corresponding quantification (F) of PHH3+ cells in control and *Srsf3* cKO embryonic hearts at E12.5. Scale bar, 50µm. Error bars indicate mean ± SEM (n = 3). Unpaired t-test with Welch's correction (**) *P* <0.01. (G) Immunofluorescence cryosection images showing expression of WT1 and tdTomato in control and *Srsf3* cKO embryonic hearts at E12.5. Scale bar, 50µm. Images representative of n = 4 embryos. *myo; myocardium, epi; epicardium.*

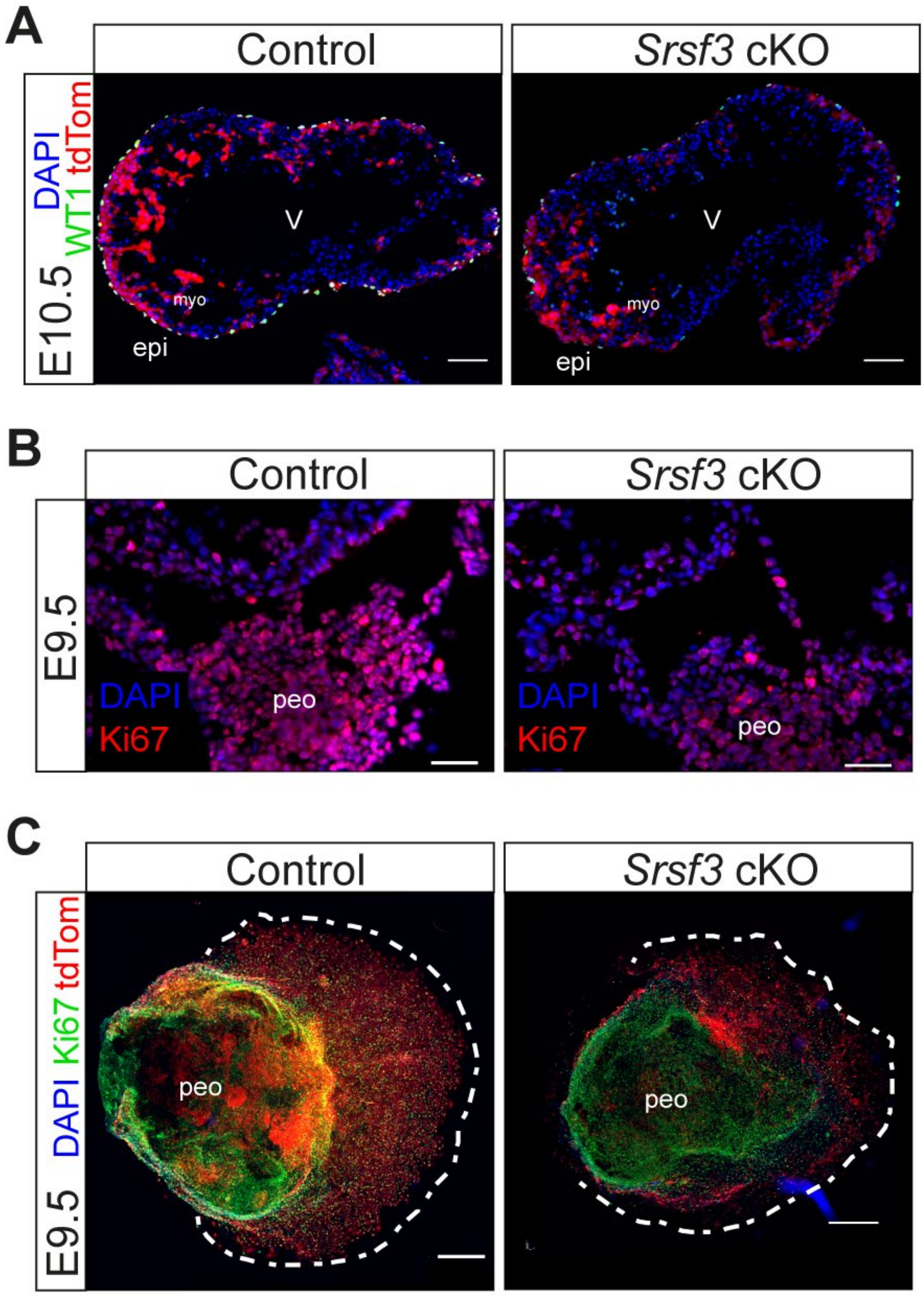

**Supplementary Figure 2. SRSF3 depletion in epicardial progenitors decreases proliferation.** (A) Immunofluorescence coronal cryosection images showing WT1+ cells on the surface of control and *Srsf3* cKO embryonic mouse hearts at E10.5. Scale bar, 100µm. Images representative of n = 3 embryos. (B) Immunofluorescence cryosection images showing expression of Ki67 in (pro)epicardial cells in control and *Srsf3* cKO embryos at E9.5. Scale bar, 50µm. Images representative of n = 3 embryos. (C) Immunofluorescence images of Ki67+ (pro)epicardial cells in control and *Srsf3* cKO E9.5 PEO explants (day 3 culture). Scale bar, 500µm. Images representative of n = 3 - 4 PEO explants. *myo; myocardium, peo; proepicardium, epi; epicardium, V; ventricle.*

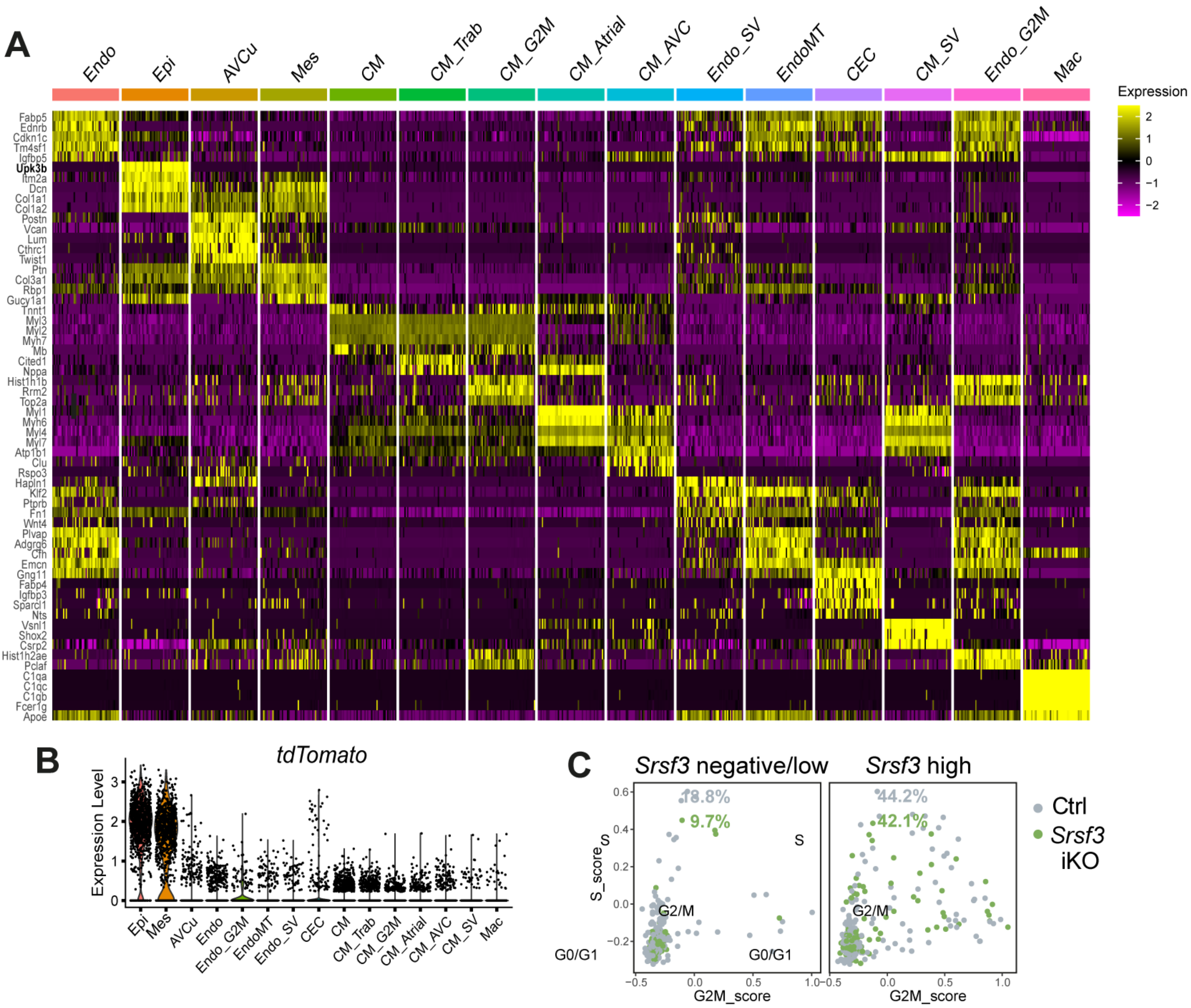

**Supplementary Figure 3. SRSF3 depletion disrupts epicardial function.** (A) Heatmap of top 5 differentially expressed genes for each cluster in the E13.5 scRNA-seq data. High expression is indicated in yellow. (B) Violin plot showing relative expression of tdTomato in individual clusters. (C) Scatter plot showing the distribution of epicardial cells in the indicated cell cycle phases when subsetted by *Srsf3* expression, in control and *Srsf3* iKO hearts. Percentage of cycling epicardial cells displayed (G1-*Mki67* positive, S and G2M phase cells). *Endo; endocardial cells, Epi; epicardium, AVCu; atrioventricular cushion, Mes; mesenchymal cells, CM; cardiomyocytes, CM_Trab; trabecular cardiomyocytes, CM_Atrial; atrial cardiomyocytes, CM_AVC; CM atrioventricular canal, Endo_SV; sinus venosus endocardial cells, EndoMT; endocardial-to-mesenchymal transition cells, CEC; coronary endothelial cells, CM_SV; sinus venosus cardiomyocytes, Mac; macrophages ,_G2M; proliferating cells.*

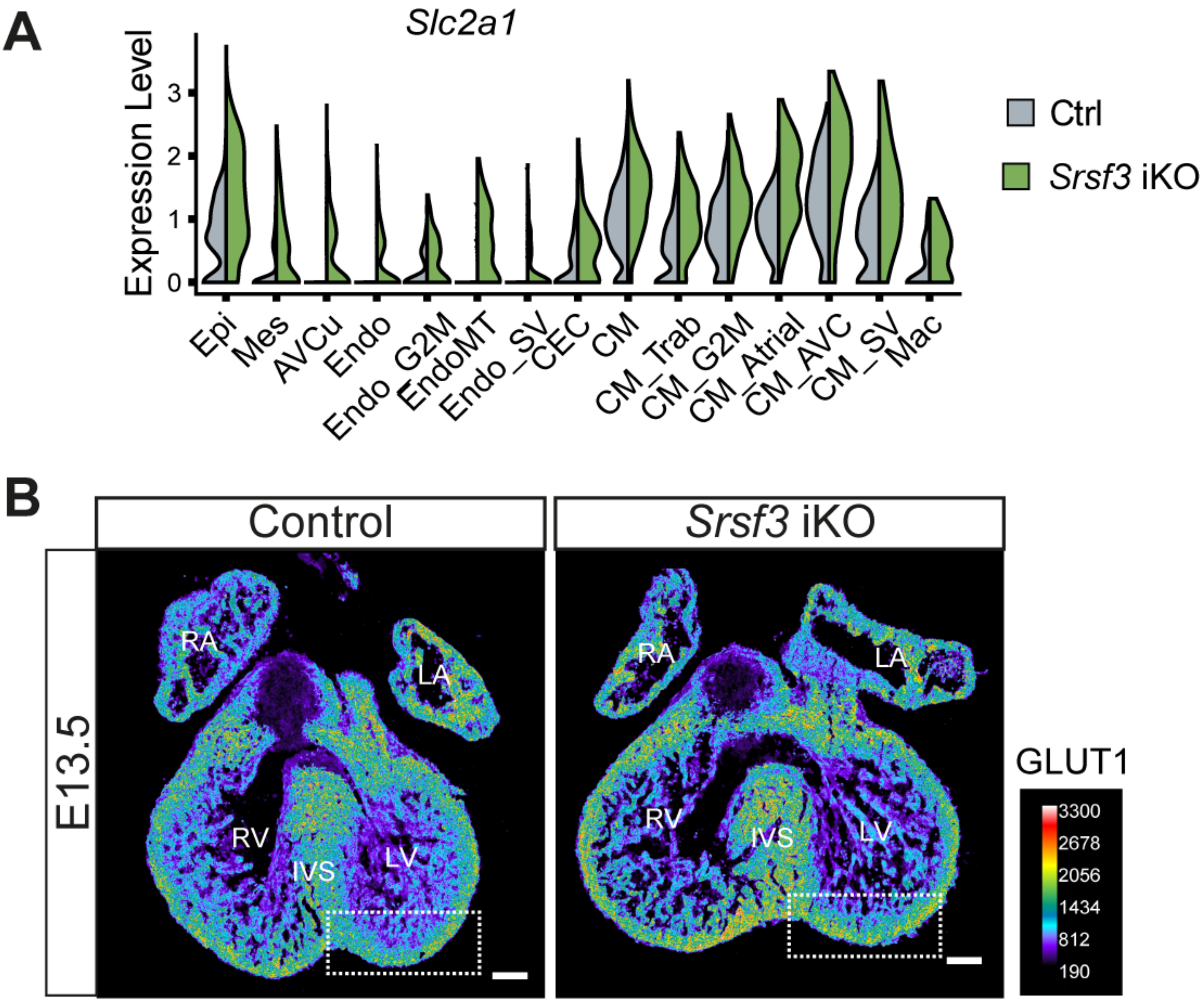

**Supplementary Figure 4. Hypoxia-responsive glucose transporter type 1 (GLUT1) is upregulated in *Srsf3* iKO hearts.** (A) Violin plot showing relative expression of *Slc2a1* in individual clusters of control and *Srsf3* iKO embryonic mouse hearts. (C) Pseudocoloured immunofluorescence images of GLUT1 expression in control and *Srsf3* iKO embryonic mouse hearts at E13.5. Scale bar, 150μm. Images representative of n = 5 hearts. *LV; left ventricle, RV; right ventricle, RA; right atrium, LA; left atrium, IVS; intraventricular septum, Endo; endocardial cells, Epi; epicardium, AVCu; atrioventricular cushion, Mes; mesenchymal cells, CM; cardiomyocytes, CM_Trab; trabecular cardiomyocytes, CM_Atrial; atrial cardiomyocytes, CM_AVC; CM atrioventricular canal, Endo_SV; sinus venosus endocardial cells, EndoMT; endocardial-to-mesenchymal transition cells, CEC; coronary endothelial cells, CM_SV; sinus venosus cardiomyocytes, Mac; macrophages , G2M; proliferating cells.*

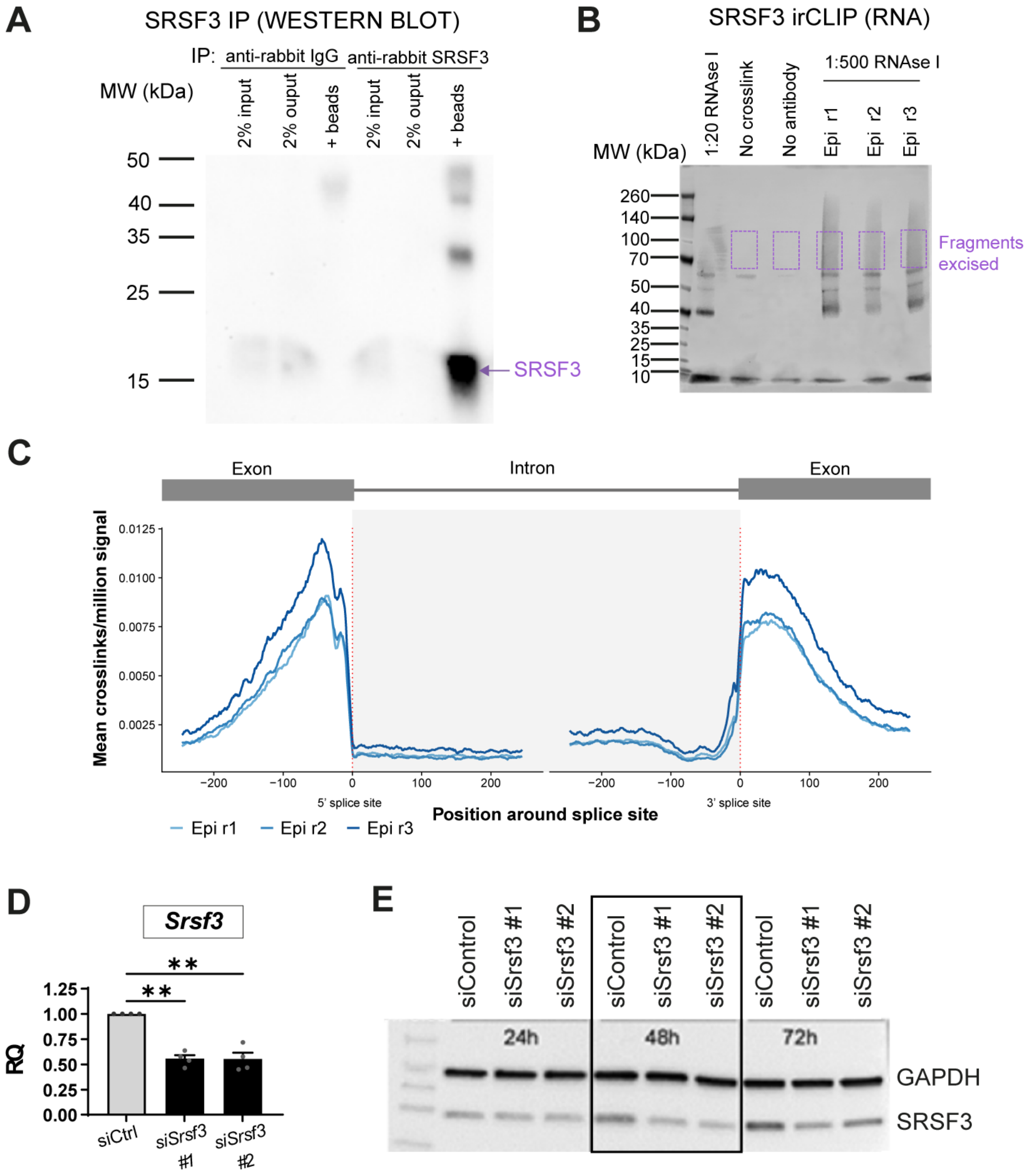

**Supplementary Figure 5. Direct SRSF3-RNA interaction in an epicardial cell line.** (A) Western blot demonstrating the relative specificity of SRSF3 immunoprecipitation. (B) Detection of SRSF3-RNA complexes; dashed boxes indicate the regions excised to purify RNA for library preparation and sequencing. (C) RNA metaprofile of the SRSF3 crosslink site distribution around 5’ and 3’ exonic splice sites. (D) qRT-PCR analysis of *Srsf3* transcript in control and *Srsf3* siRNA-transfected epicardial cell line. Error bars indicate mean ± SEM (n = 4). One-Way Welch ANOVA and Dunnett's multiple comparison test (**) $P$ <0.01. (E) Western blot of SRSF3 and GAPDH in control and *Srsf3* siRNA-transfected epicardial cell line (24h, 48h and 72h transfection). GAPDH used as loading control.

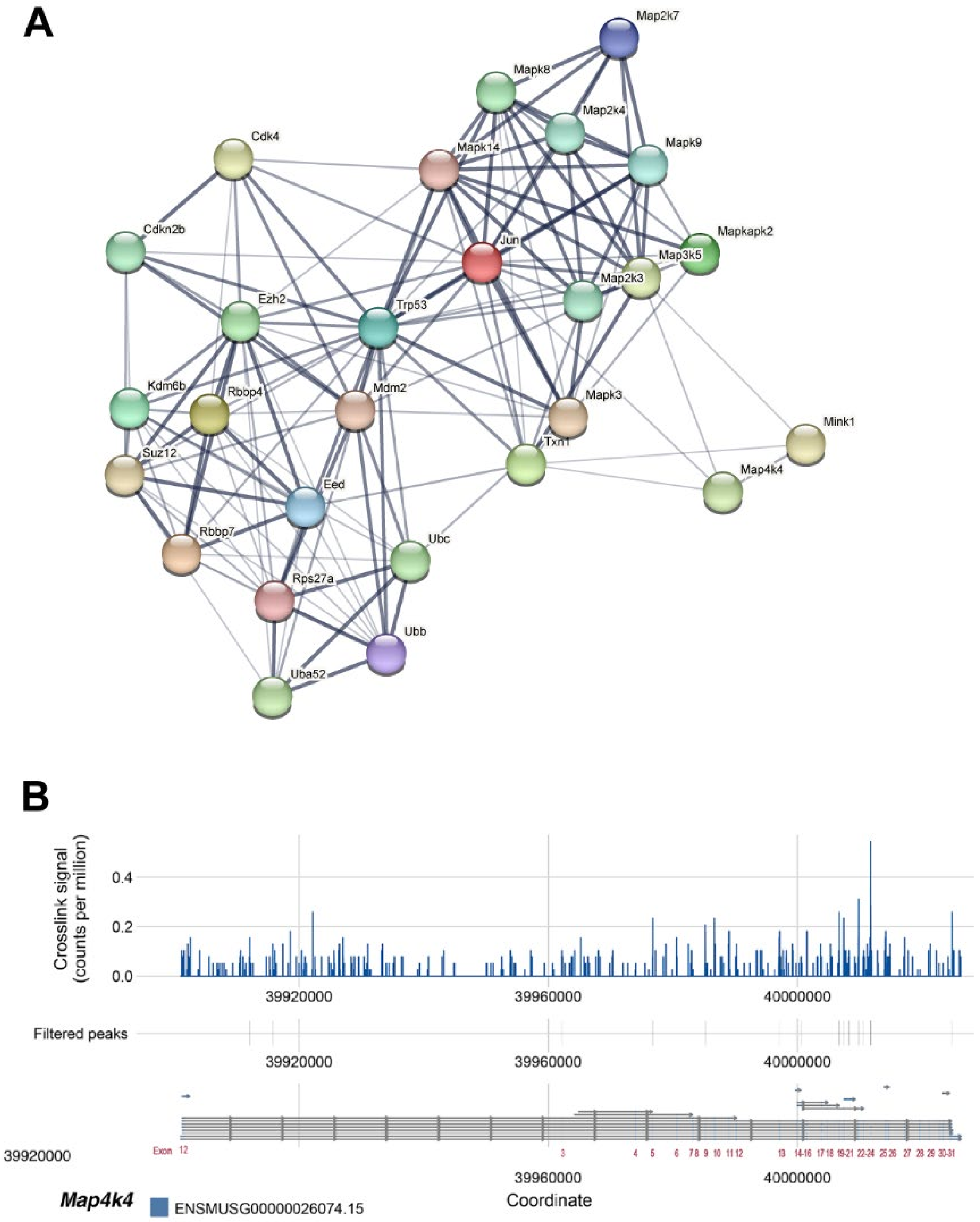

**Supplementary Figure 6. SRSF3 binds to transcripts that control cellular senescence, such as *Map4k4*.** (A) Functional association network of SRSF3 RNA targets controlling cellular proliferation and senescence in epicardial cells. (B) SRSF3 irCLIP binding profile across the 18 *Map4k4* transcripts. Significant peaks were filtered to remove those <5 nt to represent "functional" binding sites.

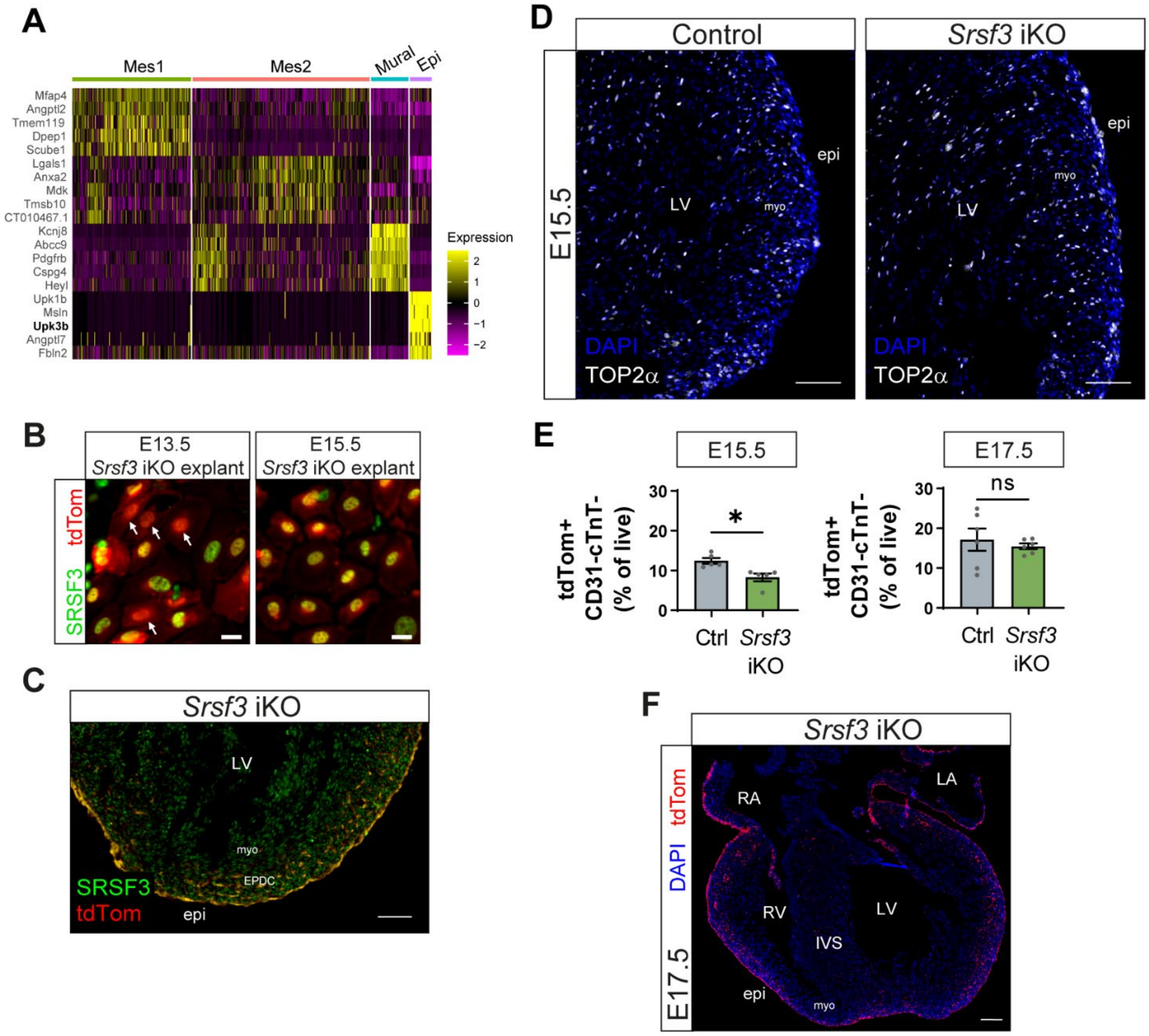

**Supplementary Figure 7. Non-recombined epicardial cells upregulate SRSF3 and hyperproliferate to compensate for the early phenotype.** (A) Heatmap of top 5 differentially expressed genes for each cluster in the E15.5 scRNA-seq data. High expression is indicated in yellow. (B) Immunofluorescence images showing expression of SRSF3 in epicardial cells of Srsf3 iKO E13.5 and E15.5 explants (day 3 culture). White arrows highlight successfully targeted SRSF3-depleted cells. Scale bar, 20μm. Images representative of n = 5 explants. (C) Immunofluorescence images showing expression of SRSF3 in the epicardium of *Srsf3* iKO hearts at E15.5. Scale bar, 100μm. Image representative of n = 4. (D) Immunofluorescence images showing expression of TOP2α in the epicardium of control and *Srsf3* iKO embryonic mouse hearts at E15.5. Scale bar, 100μm. Images representative of n = 2. (E) Flow cytometry-based quantification of percentage epicardial lineage cells (tdTomato+CD31-cTNT-) as a proportion of total live cells in control and *Srsf3* iKO embryonic mouse hearts at E15.5 and E17.5. Error bars indicate mean ± SEM (n = 5 - 6). Unpaired t-test with Welch's correction (*) *P* <0.05. (F) Fluorescence image showing epicardial lineage cells (tdTomato+) in *Srsf3* iKO embryonic mouse hearts at E17.5. Scale bar, 200μm. Images representative of n = 3 embryos. *LV; left ventricle, RV; right ventricle, LA; left atrium, RA; right atrium, IVS; intraventricular septum, myo; myocardium, epi; epicardium, EPDC; epicardium-derived cells.*

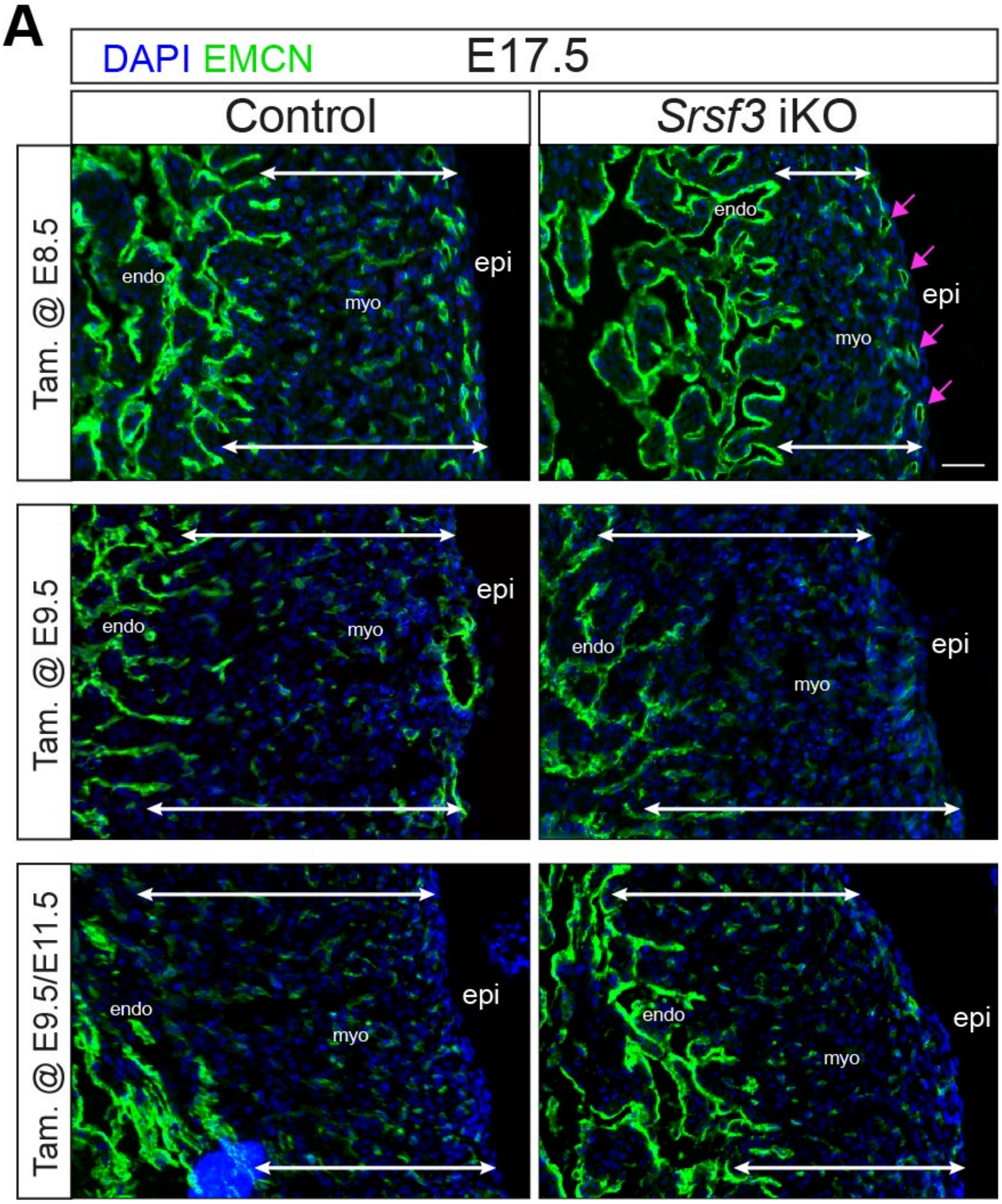

**Supplementary Figure 8. Early defects in compaction and coronary vasculature are compensated with late-stage induction of SRSF3 depletion in Srsf3 iKO hearts.** (A) Immunofluorescence cryosection images showing the endocardium and vascular network (EMCN+) in control and *Srsf3* iKO embryonic mouse hearts at E17.5. Tamoxifen induction was administered at E8.5, E9.5, and E9.5/E11.5, respectively. White arrows highlight myocardial wall thickness and magenta arrows highlight presence of superficial vessels. Scale bar, 50µm. Images representative of n = 3 - 4 embryos. *myo; myocardium, epi; epicardium, endo; endocardium.*

**Supplementary Table 1. Survival of *Tg(Gata5-Cre); Srsf3$^{fl/fl}$* embryos**

| Stage | Cre negative | Cre positive | | Litters |
|---|---|---|---|---|
| | | Het | Hom | |
| **E11.5** | 28 | 12 | 13 | 9 |
| **E12.5** | 11 | 5 | 3+1 dead | 2 |
| **E13.5** | 9 | 11 | 2 dead | 3 |
| **Born** | 10 | 6 | 0 | 4 |

*Het-heterozygote, Hom-homozygote*

### Supplementary Methods

*Immunofluorescence*

Cryosections were rehydrated, permeabilised with 0.5% Triton X-100 (PBS) for 15 mins, then incubated for 1h at RT in blocking buffer (1% bovine serum albumin (BSA) and 10% donkey serum, in 0.1% Triton X-100 PBS (PBST)). Primary antibodies (Supplemental Methods Table 1) were added, and sections incubated overnight at 4°C. Sections were washed with PBST (3 x 10 mins), then secondary antibodies were added, and incubated for 1 hour at RT, followed by more PBST washes, as above. 4',6-diamidino-2-phenylindole (DAPI) was used to stain nuclei. Cultured epicardial explants, FACS-sorted epicardial lineage and epicardial cell line[1] were subjected to the same immunofluorescence protocol, but without the initial permeabilization (0.5% Triton X-100) step. Images were acquired using Leica DM6000 fluorescence microscope or Olympus Fluoview-1000 confocal microscope and processed using Fiji software[2]. Quantification was carried out in Fiji or CellProfiler[3] software.

*Quantitative assessment of surface lumenised coronary vessels*

Whole heart images were processed using Fiji software[2]. A mask was applied based on DAPI (nuclei) staining and, through a series of dilations and shrinking, the whole cardiac tissue was segmented (region of interest 1; ROI 1). A region of 30 - 50um in depth from the epicardial border was automatically selected by segmenting 3% of the total ventricular area (outer region; ROI 2). Particle analysis was used to quantify the number, area, mean intensity, and perimeter of vessels and nuclei within the ROI 2. Particle analysis was used to quantify the area of the myocardium below the epicardium (ROI 2) and total ventricular area (ROI 1).

*Quantitative assessment of hypoxic regions in the heart*

Whole heart images were processed using Fiji software[2]. A mask was applied based on cardiac troponin staining (myocardium) and, through a series of dilations and erosions, the whole myocardium was segmented. The atria and highly trabeculated endocardial region were removed, to leave mostly compact ventricular myocardium (region of interest 1; ROI 1). Hypoxic regions with high GLUT1 levels within ROI 1 were segmented by setting a fluorescence intensity threshold (ROI 2). Particle analysis was used to quantify the total area of compact myocardium (ROI 1) and hypoxic regions (ROI 2).

*RNA extraction and qRT-PCR*

RNA was extracted from embryonic hearts using RNeasy Mini kit (Qiagen) according to the manufacturer's instructions. cDNA was prepared with High-Capacity cDNA Reverse Transcription kit (Thermo Fisher) according to the manufacturer's instructions and analysed by quantitative RT-PCR using the Fast SYBR Green Master Mix (Thermo Fisher). Primers are listed in Supplementary Methods Table 2.

*Single-cell RNA-sequencing and analysis*

For E13.5 single-cell RNA-Sequencing data, whole hearts from E13.5 embryos were dissociated as described previously[4, 5], then processed using the Chromium Single Cell 3' Reagent Kit (v3 chemistry) (10x Genomics #PN-1000092) as indicated by manufacturers' instructions. The samples were sequenced in three rounds using the NextSeq® 500/550 High Output Kit v2 (150 cycles) 400 million reads (8000 cells) (Illumina #FC-404-200) on the Illumina NextSeq 500 platform. The reads were processed using CellRanger (10x Genomics), then analysed in Seurat 3[6] as described below. Cells with more than 20% mitochondrial reads and more than 7000 genes were removed from the analysis. UMAP (Uniform Manifold Approximation and Projection) was used to visualise samples. Cell cycle S- and G2M-phase scores assigned for each individual cell (Seurat) were plotted as a scatter graph. For E13.5, three control hearts and two Srsf3 iKO hearts were used.

For the E15.5 single-cell RNA-Sequencing data, the ventricles from E15.5 embryonic hearts were dissociated as described previously[4, 5], and BD FACSAria III was used to sort live cells positive for tdTomato fluorescence into 96-well plates. The cells were processed according to the Smart-Seq2 protocol[7]. Tagmentation and library preparation was done using the Nextera XT DNA Library Prep kit and sequenced using the NextSeq 500/550 High Output Kit v2 (75 cycles) 400 million reads (Illumina, #FC-404-2005) on Illumina NextSeq 500 platform. BCL files from the sequencer were converted to FastQ with bcl2fastq (Illumina, v.2.19.1.403) using default settings. Samples with fewer reads in total than the empty wells were discarded. Reads were trimmed using default settings and "--nextera" for adapter clipping (TrimGalore v.0.4.4). Random samples were checked with FastQC (v.0.11.3) to control for the quality of the sequence. Reads were aligned to the mouse genome (GRCm38/Mm10) with STAR (v.2.3.3a) in gene counting mode with Gencode (v.M16) annotations minus the megatranscript Gm20388. Splice junctions from all samples were combined after the 1st pass alignment. Non-canonical junctions and junctions covered with < 10 reads were removed prior to running the 2nd pass alignment. The samples were further processed with Seurat. Genes with > 200 reads in at least 3 cells were retained for downstream processing. Cells were filtered to retain those with < 5% mitochondrial genes and at least 500 expressed genes. Reads were normalized to sequencing depth and scaled. For E15.5, six control hearts and four *Srsf3* iKO hearts were used.

*Whole-mount DAB staining*

Embryonic hearts were placed in 4% PFA in 1.5 ml tubes after dissection, then left O/N at 4˚C on a rotator. In between each of the following steps, hearts were washed three times for 15min in PBST. Hearts were incubated for 1h at RT in blocking buffer (5% goat serum, in PBST). An antibody against PECAM1, 1:500 dilution in PBST with 1%BSA, was added O/N at 4˚C. Hearts were incubated in biotinylated secondary antibody (Goat Anti-Armenian hamster Biotin #ab5744) 1:250 dilution in PBST for 1h. Avidin-biotin horseradish peroxidase complex (ABC Elite #Vector PK-4000) 1:50 in PBS was added for 30 min. Hearts were placed in DAB substrate (Vector peroxidase substrate kit #SK-4100). When vessels were visible, the reaction was stopped by rinsing the hearts with ddH2O. The hearts were imaged on a stereo microscope.

*Western Blotting*

Protein was extracted using RIPA buffer (50 mM Tris-HCl pH 7.4, 1% Triton X-100, 0.1% SDS, 150 mM NaCl, 2 mM EDTA) with Protease Inhibitors (Sigma #P8340) and Phosphatase Inhibitors (Sigma #P5726). Protein concentration was quantified using Pierce BCA Protein Assay kit (ThermoScientific #23227), then 6 µg of protein was loaded on a 4-20% MiniProtean Gel (BioRad #4561096). The samples were transferred using the Trans-Blot Turbo Transfer Pack (BioRad #1704456), blocked in 8% skimmed milk powder/ TBST (20 mM Tris, pH 7.5, 150mM NaCl, 0.1% Tween 20) for 1h, then incubated in 1:1000 SRSF3 (Abcam #ab73891) and 1:5000 GAPDH (Abcam #ab8245) in blocking buffer (5% skimmed milk powder/TBST) O/N at 4°C, followed by 3 washes of 5 min each in TBST and incubation with HRP-conjugated secondary antibodies, diluted 1:2500 in blocking buffer. The blot was rinsed 3 times for 5 min in TBST, then the samples were detected using ECL™ Western Blotting Detection Reagent (Sigma #GERPN2209).

*Flow Cytometry*

For quantifying the epicardial lineage, E15.5 or E17.5 whole hearts were dissociated, and surface stained with viability dye, then pre-treated for 5 min on ice with TruStain FcX antibody to block Fcγ receptors. Cells were subsequently stained with BV605 rat anti-CD31 antibody, on ice for 0.5hr. Cells were fixed and permeabilised, then intracellularly stained with cTNT as previously described[4, 5]. Cells were stored in IC fixation buffer (Thermo Fisher, #00-8222-49) until acquisition.

*Statistical analysis*

Statistical analysis was performed in GraphPad Prism 9 software. Unpaired t-test with Welch's correction was used to compare two experimental groups. One-way Welch ANOVA and

Dunnett's multiple comparison test was used to compare more than two experimental groups. P value lower than 0.05 was considered significant. Details for statistical analyses, including biological replicate numbers, are included in figure legends.

*Supplementary Methods Table 1.* ***List of antibodies used for Immunostaining and Flow Cytometry***

| **Antibody** | **Company** | **Catalogue No.** | **Species** | **Dilution** |
|---|---|---|---|---|
| WT1 | Abcam | Ab89901 | Rabbit | 1:100 |
| SRSF3 | Abcam<br>Millipore | Ab73891<br>MABE116 | Rabbit<br>Mouse | 1:200<br>1:100, ICC |
| PECAM1 | Abcam | Ab119341 | Ar. Hamster | 1:100 |
| Ki67 | eBioscience | Ab569882 | Rat | 1:100 |
| CC3 | Cell Signaling Technology | 9664 | Rabbit | 1:100 |
| PHH3 | Santa Cruz | Sc-8656 | Rabbit | 1:100 |
| α-s-actinin | Abcam | Ab9465 | Mouse | 1:100 |
| CCND1 | Abcam | Ab16663 | Rabbit | 1:100 |
| EMCN | Santa Cruz | Sc-65495 | Rat | 1:100 |
| TOP2α | Abcam | ab52934 | Rabbit | 1:100 |
| GLUT1 | Abcam | Ab115730 | Rabbit | 1:100 |
| BV421 cTNT | BD Biosciences | 565618 | Mouse | 1:200 |
| AF657 cTNT | BD Biosciences | 565744 | Mouse | 1:100, IF |
| BV605 CD31 | Biolegend | 102427 | Rat | 1:50 |
| TruStain FcX | Biolegend | 101319 | Rat | 1:50 |
| Zombie Aqua Fixable Viability Kit | Biolegend | 423101 | N/A | 1:1000 |

*Supplementary Methods Table 2.* ***qRT-PCR primers used***

| Gene | Forward ( 5' -> 3') | Reverse ( 5' -> 3') |
|---|---|---|
| *Srsf3* | GCGCAGGTACTTGAGAGA | ATCGGCTACGAGACCTAGAGA |
| *Actb* | CTGTCGAGTCGCGTCCACC | GCTTTGCACATGCCGGAGC |
| *Canx* | ATGGAAGGGAAGTGGTTACTGT | GCTTTGTAGGTGACCTTTGGAG |
| *Ywhaz* | GAAAAGTTCTTGATCCCCAATGC | TGTGACTGGTCCACAATTCCTT |
| *Ccnd1* | GTTCATTTCCAACCCACCCTC | AGAAAGTGCGTTGTGCGGTAG |
| *Map4k4* | CATCTCCAGGGAAATCCTCAGG | TTCTGTAGTCGTAAGTGGCGTCTG |
| *Map4k4 IE16/17* | GAGCCACCCGTGCCTTCCC | GACTGAGCCCCTCACTGCG |
| *Map4k4 SE16/17* | CGCCCCCGCAGCAGCAGGA | GTTCTCACGGAACCTCCCGAGCT |
| *Map4k4 SE16* | CGCCCCCGCAGCAGCAGGA | ACCACTGTACCTCCCGAGC |
| *Map4k4 SE17* | CGCCCCCGCAGCAGCAGGA | GTTCTCACGGAACCTGTGGCTCC |

*Supplementary Methods Table 3.* ***RNAscope probes and TSA plus fluorophore***

| **Probe** | **Catalogue No.** |
|---|---|
| Ndrg1 | 461971 |
| dapB (negative control) | 320878 |
| 3-plex Negative control | 320871 |
| **TSA fluorophore** | **Catalogue No.** |
| TSA plus Cy3/Cy5 1:1000/1:1500 | NEL753001KT |
